# The lactonase BxdA mediates metabolic adaptation of maize root bacteria to benzoxazinoids

**DOI:** 10.1101/2023.09.22.559061

**Authors:** Lisa Thoenen, Marco Kreuzer, Matilde Florean, Pierre Mateo, Tobias Züst, Caitlin Giroud, Liza Rouyer, Valentin Gfeller, Matheus D. Notter, Eva Knoch, Siegfried Hapfelmeier, Claude Becker, Niklas Schandry, Christelle A. M. Robert, Tobias G. Köllner, Rémy Bruggmann, Matthias Erb, Klaus Schlaeppi

## Abstract

Root exudates contain secondary metabolites that affect the plant’s root microbiome. How microbes cope with these bioactive compounds, and how this ability shapes root microbiomes remain largely unknown. We investigated how maize root bacteria metabolise benzoxazinoids, the main specialised metabolites of maize. Diverse and abundant bacteria metabolised the major compound (6-methoxy-benzoxazolin-2-one, MBOA) in the maize rhizosphere to 2-amino-7-methoxyphenoxazin-3-one (AMPO). By contrast, bacteria isolated from Arabidopsis, which does not produce benzoxazinoids, were unable to metabolise MBOA. Among *Microbacteria* strains, this differential metabolisation allowed to identify a conserved gene cluster containing the lactonase *bxdA*. BxdA converts MBOA to AMPO in vitro and we show that this capacity provided bacteria a growth benefit under carbon-limiting conditions. Together these results reveal that maize root bacteria - through BxdA - are metabolically adapted to the benzoxazinoids of their host. We propose that metabolic adaptation to plant-specialised compounds shapes root bacterial communities across the plant kingdom.

## Introduction

Plant microbiomes fulfil key functions for plant and ecosystem health. Root-associated microbes promote plant growth, provide nutrients, and protect plants from pathogens^1,2^. While some root microbes are ubiquitous, many microbes form specific relationships with their host plants, and host plants often exert substantial control over the structure and function of their microbiome. Plants primarily shape their root-associated microbiome through the secretion of root exudates, which can account for up to one-fifth of the plant’s assimilated carbon^3^. Root exudates may attract, nourish, or repel soil microbes and contain primary metabolites including sugars, amino acids, organic acids and fatty acids, as well as secondary metabolites. The latter, also called specialised metabolites, govern the plant’s interactions with the environment, and among other functions, they increase biotic and abiotic stress tolerance^4^. A key function of exuded specialised metabolites is to shape the root microbiomes^5–7^, documented with examples including glucosinolates, camalexins, triterpenes, and coumarins from *Arabidopsis thaliana*^5^, the saponin tomatine from tomato^8^, and benzoxazinoids^9–13^, diterpenoids^14^, zealexins^15^ and flavonoids^16^ from maize.

Benzoxazinoids are multifunctional indole-derived metabolites produced by *Poaceae*, including crops such as wheat, maize, and rye^17^. These compounds accumulate in leaves as chemical defences against insect pests and pathogens^17^ and are exuded from the roots as phytosiderophores^18^ and antimicrobials^19–21^. Benzoxazinoids directly shape the root and rhizosphere microbiomes^9–11,22^, and when metabolised to aminophenoxazinones by soil microbes, they also become allelopathic, inhibiting the germination and growth of neighbouring plants^17^. DIMBOA-Glc is the main root-exuded benzoxazinoid of maize^11^, and its chemical fate in soil is well understood. In Fig. S1 we document full names, structures and relatedness of all compounds relevant to this study. Upon exudation, plant- or microbe-derived glucosidases^17^ cleave off the glucose moiety to form DIMBOA, which spontaneously converts to more stable MBOA^23^. In soil, MBOA has a half-life of several days and can be further metabolised to reactive aminophenols by microbes^17^. Three routes to different metabolite classes are known: route (I), favoured under aerobic conditions^24^, forms aminophenoxazinones such as AMPO and AAMPO; route (II) results in acetamides such as HMPAA through acetylation^25^, or alternatively, route (III) yields malonic acids such as HMPMA through acylation^25^. Route I is certainly relevant for the rhizosphere as the AMPO can be detected in soils of cereal fields over several months^23^. While the chemical pathways of benzoxazinoid metabolisation are well-defined, the responsible microbes and enzymes remain largely unknown (but see below).

Benzoxazinoids and their metabolisation products have antimicrobial properties. Yet, it remains poorly understood, how microbes cope with these bioactive plant metabolites^19–21,26,27^. We discriminate metabolite-microbe interactions as ‘native’ or ‘non-host’, the latter referring to context where root microbes and root metabolites do not originate from and occur in the same host. Recently, we demonstrated that ‘native’ root bacteria (isolated from maize) tolerated the maize-originating benzoxazinoids better compared to ‘non-host’ bacteria isolated from Arabidopsis^19^. This suggested that native maize bacteria were adapted and have evolved strategies to tolerate these compounds. Evolution of tolerance could either involve reduced sensitivity of molecular targets in the bacteria or improved and/or specialised strategies for metabolic detoxification. Adapted bacteria may metabolise plant-derived compounds either by conversion to less toxic compounds, or by degrading them entirely. Metabolisation of plant-derived compounds may not only reduce toxicity but also have added benefits for bacterial growth. *Pseudomonas* or *Sphingobium* bacteria for instance use exuded triterpenes or tomatine as carbon sources, respectively^28,8^. These examples suggest that native bacteria have evolved specialized adaptations to metabolise secondary metabolites in root exudates of their host – this hypothesis remains untested.

Several soil microbes have been found to metabolise benzoxazinoids. Examples of compound conversions include APO formation from BOA (non-methoxylated form of MBOA, Fig. S1) by *Acinetobacter* bacteria^29^, formation of the acetamide HPAA from BOA by the fungus *Fusarium sambucus*^25^, or accumulation of APO from BOA upon co-culture of *Fusarium verticillioides* with a *Bacillus* bacterium^30^. Testing different soil microbes from various environments revealed that they differed strongly in their metabolic activities but that degradation resulted in the expected sequence of compounds from DI(M)BOA-Glc to DI(M)BOA to (M)BOA^13^. First insights into the molecular mechanisms include the identification of the metal-dependent hydrolase CbaA from *Pigmentiphaga* bacteria that degrade modified benzoxazinoids^31^, and of a metallo-β-lactamase (MBL1) from the maize seed endophytic fungus *Fusarium verticilloides* that degrades BOA to the malonamic acid HPMA^32^. Benzoxazinoid metabolisation by microbes has commonly been studied with diverse microbes isolated from different soil environments. Microbial metabolisation of benzoxazinoids and its genetic basis have not yet been investigated in the native context of root microbes from benzoxazinoid-exuding plants.

To uncover ecological context and biochemistry of microbial benzoxazinoid metabolisation, we systematically screened native maize and non-host Arabidopsis root bacteria. Using metabolite analyses, genetics, comparative genomics, and biochemical validation, we characterised benzoxazinoid-metabolising maize root bacteria and identified the underlying genetic mechanisms. We found a conserved gene cluster for benzoxazinoid metabolisation with a lactonase that catalyses the degradation of MBOA. Our work thereby uncovered a metabolic adaption of root bacteria to host-exuded specialised metabolites.

## Results

### Taxonomically widespread and abundant maize root bacteria form AMPO

Screening maize root bacteria (i.e., the ‘MRB collection’) for MBOA tolerance^19^, we observed that some liquid cultures, including *Sphingobium* LSP13 and *Microbacterium* LMB2, turned red (Fig. S2). Analysis of the liquid media by UPLC-MS revealed that these bacteria degraded MBOA, and NMR analysis with comparison to an analytical standard confirmed the formation of AMPO, which has a dark red colour. The colour change to red also manifests on MBOA-containing agar plates (Fig. S3) and thus, served as visual screen for AMPO-formation in subsequent experiments.

To define the distribution of AMPO-forming bacteria, we screened our culture collection^19^ on MBOA-containing plates and classified the strains as *non*, *weak* or *strong* AMPO formers (see Fig. S3). We identified 44/151 strains belonging to six genera from two phyla with colour changes to light or dark red (Fig. 1a). Strong AMPO-formers were strains from *Microbacteria* (17/28 tested) and *Pseudoarthrobacter* (3/3), both of the phylum Actinobacteriota. Among Pseudomonadota, *Sphingobium* (13/13) and *Enterobacter* (4/4) strains were strong AMPO-formers too, while *Rhizobium* (6/7) and *Acinetobacter* (1/1) isolates were weak AMPO-formers. Metabolite profiling confirmed AMPO-formation in liquid cultures with MBOA (see below). We concluded that AMPO-formation is a taxonomically widespread trait among maize root bacteria.

**Figure 1:**
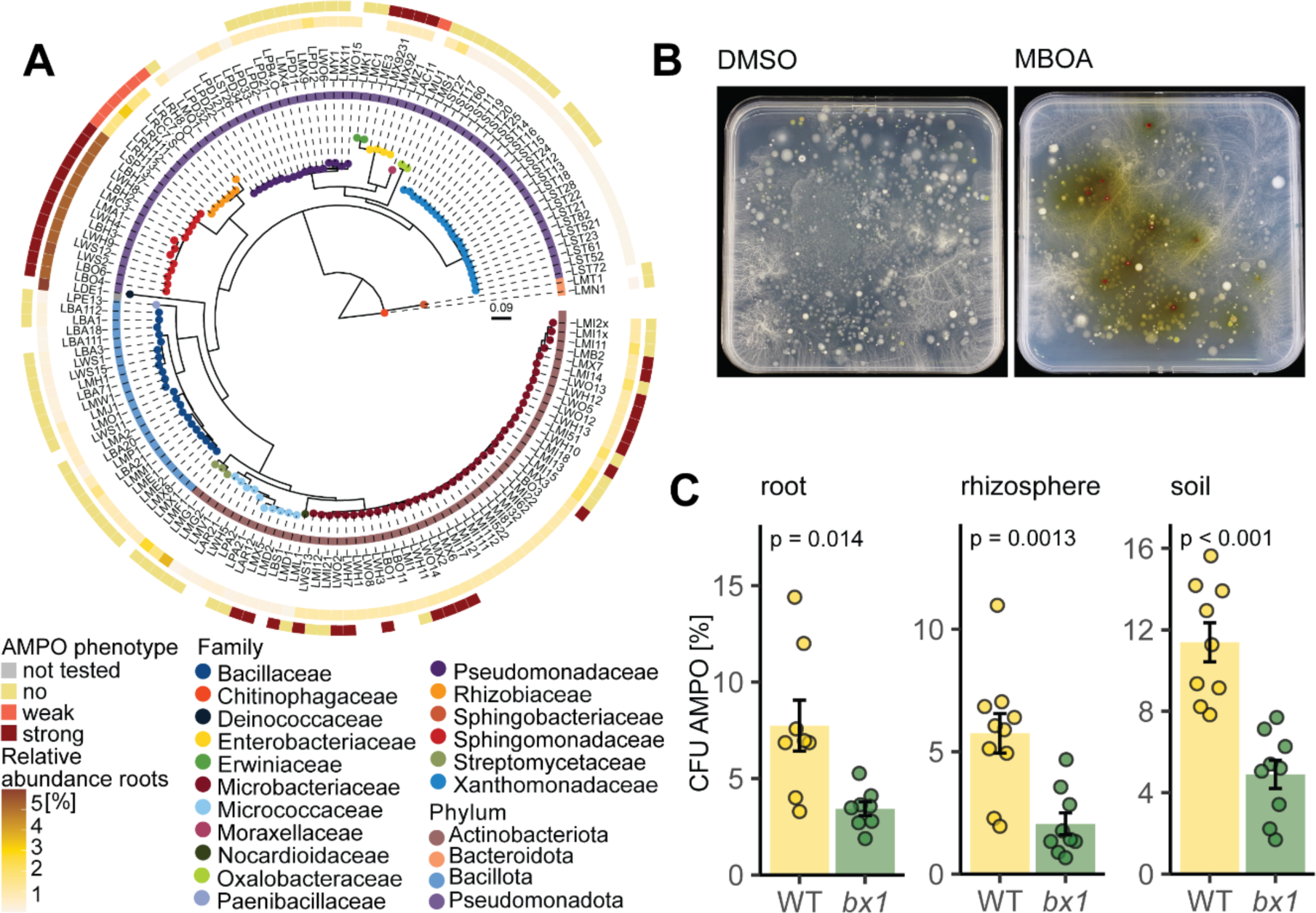
AMPO-forming colonies are abundant on benzoxazinoid-exuding maize roots. **A)** Maximum likelihood phylogeny, constructed from the alignment of 16S rRNA gene sequences of maize root bacteria Leaf nodes are coloured by family taxonomy and the ring next to the strain IDs reports phylum taxonomy. The inner ring is coloured according to the relative abundance (%) of the corresponding sequence in the microbiome profile of the roots, from which the isolates were isolated. The outer ring displays the phenotype of the strains in the plate assay classified as “strong AMPO-former” based on a strong red colouring of the agar plate, “weak AMPO-former” for strains colouring the media to lighter red or “no AMPO-former” for strains not showing a colour change compared to the control root extracts plated on bacterial growth medium supplemented with DMSO (left) and MBOA (right) grown for 10 days. AMPO-forming colonies appear red on the MBOA-supplemented medium. **C)** Percentage of total colony forming units (CFU) that form AMPO on wild-type (WT) or benzoxazinoid-deficient *bx1* mutant roots, in rhizosphere and soil. Means ± SE bar graphs and individual data points are shown (WT n = 8, *bx1* n = 9). Results of pairwise t-tests are shown inside the panels.

To approximate the abundance of AMPO-forming bacteria in microbiomes, we mapped the identified strains to maize root microbiome datasets. First, mapping them to roots from which they were isolated from^11^, revealed that they accounted for 9% of the community, with *Sphingobium* contributing most (5.3%, strong AMPO-former), followed by *Rhizobium* (1.6%, weak), *Microbacteria* (1%, strong) and *Enterobacter* (0.7%, strong; Fig. 1a). Second, in maize root microbiomes from field data^9^, community abundances ranged from 2.9% (Changins, CH), to 6.7% (Aurora, US) and 14.9% (Reckenholz, CH; Fig. S4). Then, to confirm the abundance of AMPO-formation in natural microbial communities, we plated extracts of maize roots, rhizospheres, and soil of a pot experiment with wild-type and benzoxazinoid-deficient *bx1* mutant plants on MBOA-containing agar plates and determined the proportion of red colonies (Fig. 1b). In extracts from wild-type plants, ∼7.7 % of the root bacteria, ∼5.8% of the rhizosphere bacteria and ∼11.4% of the soil bacteria formed AMPO (Fig. 1c). In extracts from *bx1* mutants, the proportion of AMPO-forming bacteria decreased by more than 50%. Together with the mapping, these results suggest that AMPO-forming bacteria are abundant and enriched by benzoxazinoids in root microbiomes of maize.

### AMPO-formation is specific for maize root bacteria

Using the same plate assay, we tested for AMPO-formation among root bacteria from different host plants. We compared root extracts from maize with wheat (*Triticum aestivum*), which accumulate less and predominantly non-methoxylated benzoxazinoids (i.e., BOA instead of MBOA)^33–35^, and with lucerne (*Medicago sativa*), oilseed rape (*Brassica napus*) and Arabidopsis (Fig. 2a), all of which do not produce benzoxazinoids. We found the highest proportion of AMPO-forming colonies on maize roots (∼7.7 %), followed by *Brassica* (∼1 %), *Triticum* (∼0.5 %), *Medicago* (∼0.07 %) and Arabidopsis (∼0.002 %). These findings highlight that AMPO-forming bacteria are specifically enriched on roots of maize plants.

**Figure 2:**
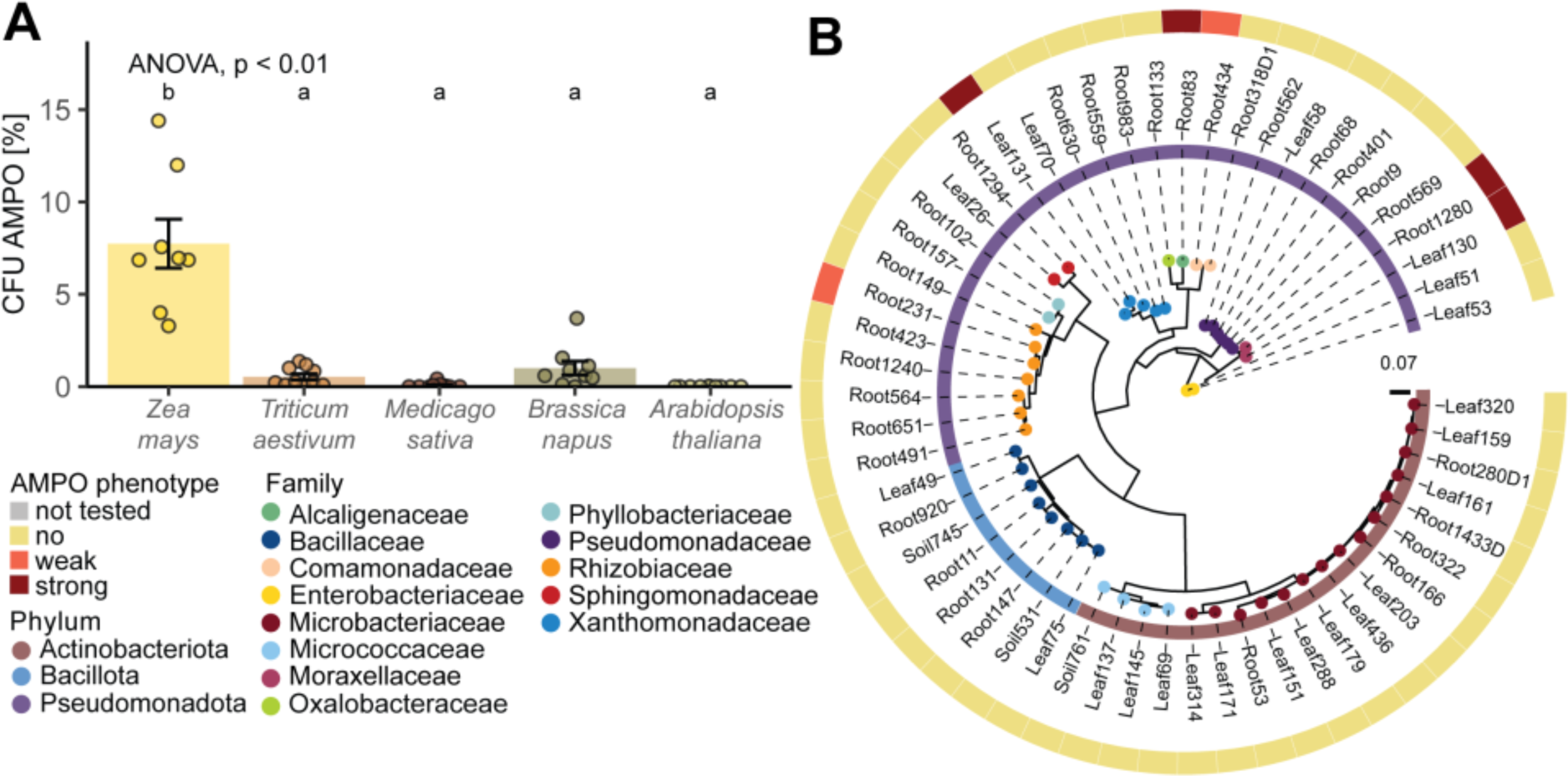
AMPO-formation in root bacteria from other host plants. **A)** Percentage of colony forming units (CFU) of AMPO-forming colonies in root extracts of benzoxazinoid producing plants *Zea mays* (maize), *Triticum aestivum* (wheat) and non-benzoxazinoid producing plants *Medicago sativa* (lucerne), *Brassica napus* (oilseed rape) and *Arabidopsis thaliana*. Means ± SE and individual data points are shown (n = 10, except maize n = 8) ANOVA and compact letter display of all pair-wise comparisons (Significance-level: FDR-corrected p < 0.05) of estimated marginal means are shown. **B)** Maximum likelihood phylogeny, constructed from the alignment of 16S rRNA gene sequences of Arabidopsis bacteria (AtSphere). Leaf nodes are coloured by family taxonomy and the ring next to the strain IDs reports phylum taxonomy. The ring displays the phenotype of the strains in the plate assay classified as “strong AMPO-former” based on a strong red colouring of the agar plate, “weak AMPO-former” for strains colouring the media to lighter red or “no AMPO-former” for strains not showing a colour change compared to the control.

To confirm this finding, we screened a collection of Arabidopsis bacteria^36^ for AMPO-formation using the classification approach. On MBOA-containing plates 2/57 strains classified as weak and 4/57 as strong AMPO-formers (Fig. 2b). A subset of Arabidopsis bacteria was further tested in liquid culture alongside two strong and two weak AMPO-formers from maize (Fig. S5a). None of the Arabidopsis strains efficiently degraded MBOA compared to the strong AMPO formers of maize. Only *Acinetobacter* (Root1280 and Leaf130) and *Variovorax* (Root434) formed low amounts of AMPO. This screening of Arabidopsis bacteria confirmed the results from plating root extracts (Fig. 2a) and revealed that efficient AMPO-formation is a specific trait of bacteria isolated from benzoxazinoid-exuding maize roots.

### Strong MBOA-degradation is required for AMPO-formation

For chemical validation of the AMPO formers and to investigate whether maize root bacteria also degrade MBOA without forming AMPO, we exposed 50 strains to 500 μM MBOA in liquid cultures and quantified MBOA metabolisation using UPLC-MS. Because the metabolite data (Fig. S5B) is not normalized by bacterial growth, we report qualitative classifications in Fig. 3a. We classified 30/50 strains not to degrade MBOA (±10% of the control), while 14 strains partially degraded MBOA (‘weak MBOA-degraders’; >30% degraded compared to the control) and 6 strains were ‘strong MBOA-degraders’ (>90% degraded). 8/50 strains were classified as ‘strong AMPO-formers’, 9 strains forming lower amounts of AMPO (‘weak AMPO-formers’, <10% of max. AMPO-former) while most strains were non-AMPO formers (<0.1% of max. AMPO-former) and we noticed 5 strains that metabolised AMPO further to AAMPO. We also noticed that the initial amount of MBOA disappeared in the cultures of the strong MBOA-degraders while low amounts of AMPO formed (Fig. S5B). This screening allowed the following conclusions: the liquid assay confirmed the AMPO-formers previously classified on plates (Fig. 1a), many bacteria degraded MBOA without forming AMPO, which is consistent with the existence of alternative MBOA degradation pathways (Fig. S1), and importantly, strong MBOA-degraders were strong AMPO-formers. Exceptions were only *Enterobacter* LME3 and *Paenarthrobacter* LAR21, which formed (A)AMPO without a strong decrease in MBOA, suggesting the existence of multiple ways to form AMPO from MBOA.

**Figure 3:**
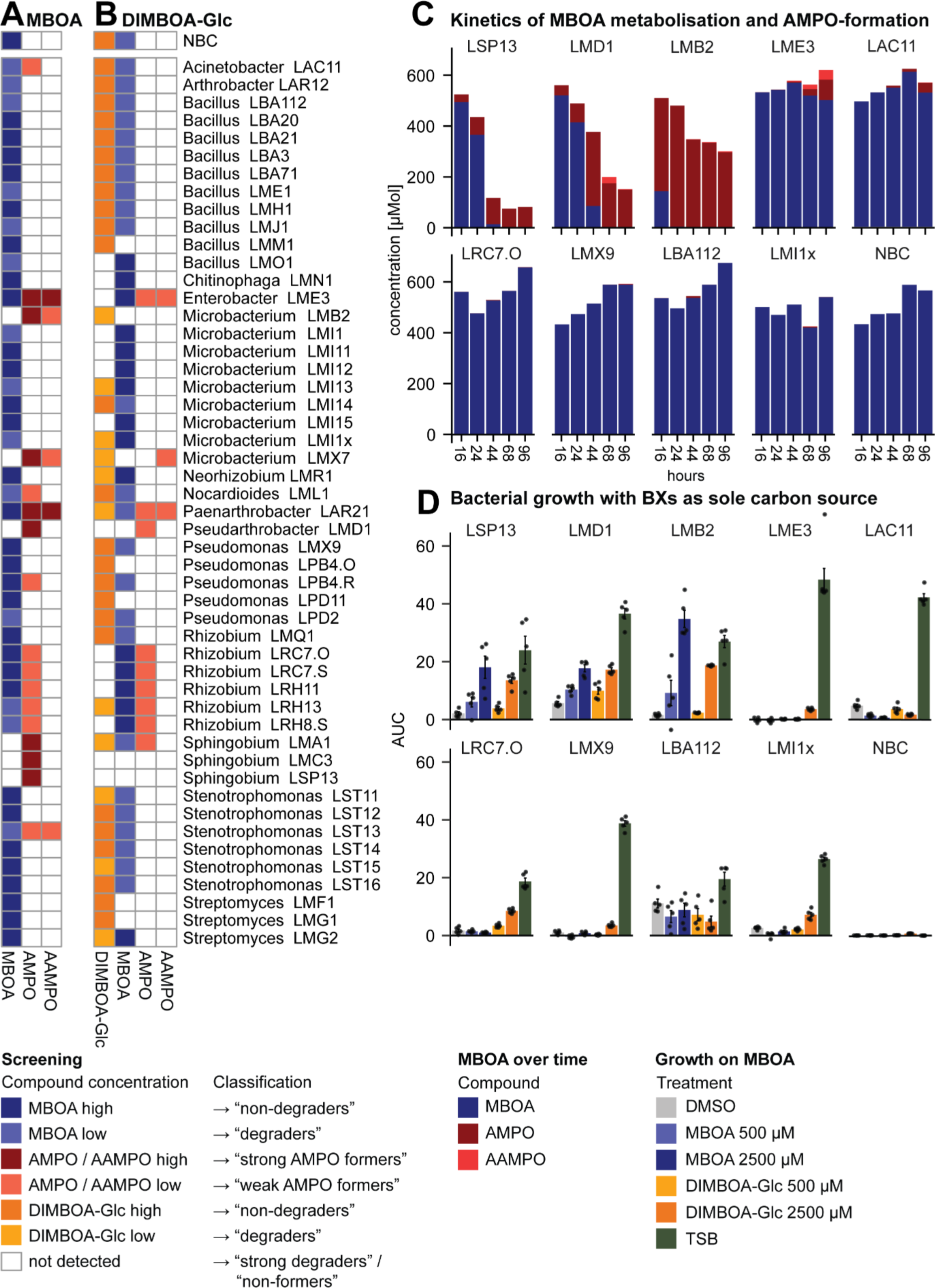
Metabolisation of benzoxazinoids and use as sole carbon source by maize root bacteria. **A)** Heatmap displaying qualitative classifications for MBOA and its metabolisation products AMPO and AAMPO and for **B)** DIMBOA-Glc and its metabolisation products MBOA, AMPO and AAMPO in liquid cultures of 50 tested maize root bacteria. “No bacteria control” NBC only contains the medium supplemented with the respective chemicals. **C)** Metabolisation of MBOA to AMPO and AAMPO over time (16h, 24h, 44h, 68h, 96h) for selected single strains: strong AMPO-formers *Sphingobium* LSP13, *Pseudoarthrobacter* LMD1, *Microbacterium* LMB2, *Enterobacter* LME3 and weak AMPO-formers *Acinetobacter* LAC11 and *Rhizobium* LRC7.O and *Pseudomonas* LMX9, *Bacillus* LBA112 and *Microbacterium* LMI1x as negative controls. All measurements were made from three independently grown cultures, which were pooled in equal ratios before metabolite analysis. **D)** Testing MBOA as sole carbon source reported as bacterial growth during 68 h (area under the curve, AUC) of the same single strains in minimal medium supplemented with DMSO (negative control), MBOA, or DIMBOA-Glc each in two concentrations (500 µM or 2’500 µM) and in TSB as positive growth control. Means in bar graphs and individual data points are shown (n = 5).

Since maize bacteria are not first exposed to MBOA on roots, we analysed metabolisation of DIMBOA-Glc, the main compound in maize exudates. Of note, maize exudates also contain DIMBOA (Fig. S1), we did not analyse it because of spontaneous conversion to MBOA in absence of bacteria in the assay (Fig. S5C). DIMBOA-Glc is commercially not available, thus we purified it from maize (traces of other co-purified benzoxazinoids, Fig. S5C). Half of the strains were classified as ‘non-degraders’ of DIMBOA-Glc (±10% of the control; Fig. 3b). The DIMBOA-Glc metabolising strains were classified as ‘weak degraders’ (11 strains; >30% degraded compared to control) and ‘strong degraders’ (14 strains, >90% degraded). These DIMBOA-Glc degrading strains generally accumulated MBOA in their cultures, while only a few strains subsequently formed low amounts of (A)AMPO. Importantly, strong degraders of DIMBOA-Glc were not necessarily strong MBOA-degraders, revealing that these are two uncoupled traits in maize root bacteria.

To characterize the kinetics of MBOA-degradation and AMPO-formation, we performed a time-series experiment with four strong (*Sphingobium* LSP13, *Pseudoarthrobacter* LMD1, *Microbacterium* LMB2, and *Enterobacter* LME3) and two weak AMPO-formers (*Acinetobacter* LAC11 and *Rhizobium* LRC7.O) alongside three non-AMPO formers (*Pseudomonas* LMX9, *Bacillus* LBA112 and *Microbacterium* LMI1x). Rapid and strong AMPO-formation was coupled with a strong decrease of MBOA (LSP13, LMD1 and LMB2) while low amounts of AMPO formed with time and without much decrease of MBOA (LME3 and LAC11; Fig. 3c). Neither MBOA-degradation nor AMPO-formation was detected in LRC7.O and the negative controls. Together with Fig. 3a these experiments indicate at least two ways to form AMPO from MBOA: (i) AMPO is formed slowly and most likely as the only product from MBOA or (ii) AMPO is rapidly formed in course of a fast and strong degradation of MBOA.

Literature suggests that MBOA degrades to the reactive intermediate AMP (Fig. S1), of which two molecules spontaneously form AMPO in the presence of oxygen^24^. To confirm the requirement for oxygen in AMPO formation, we cultivated the bacteria LMB2, LMD1 and LSP13 both under aerobic and anaerobic conditions. Much less MBOA was degraded and much less AMPO formed in anaerobic conditions (Fig. S6). While this result should be interpreted with caution, as bacterial growth was also reduced in absence of oxygen, it is in line with spontaneous AMPO formation from AMP in presence of oxygen^24^.

### AMPO-formation benefits bacterial growth

Because the degradations of DIMBOA-Glc and MBOA are key steps to form AMPO, (Fig. 3ab), we tested whether these metabolic traits provided AMPO-forming strains a growth benefit. We grew the same strains of the kinetic analysis (Fig. 3c) in minimal media with DIMBOA-Glc or MBOA as sole carbon source. All strains grew well in the positive growth control with TSB medium and the non-degraders of both DIMBOA-Glc and MBOA (LAC11 and LMX9, LBA112) did not grow on either carbon source (Fig. 3d). LME3, LRC7.O and LMI1x, being capable to degrade DIMBOA-Glc but not MBOA, partially benefited in minimal medium with the high concentration of DIMBOA-Glc but not MBOA. In contrast, the strong DIMBOA-Glc- and MBOA-degraders (LSP13, LMD1, LMB2) strongly increased their cell numbers compared to the controls. Together, these results reveal that the capacities to degrade DIMBOA-Glc and MBOA are directly associated with growth benefits under carbon-limiting conditions.

### AMPO-formation varies within *Microbacteria*

To identify the genetic basis of AMPO-formation, we took advantage of the phenotypic diversity in *Microbacteria* (Fig. 1a) and chose all isolates from maize^19^ (n=18), Arabidopsis^36^ (n=17), and other plants we had available in the laboratory (n=4, Dataset S1; see methods). We tested this set of 39 *Microbacteria* for AMPO-formation using the plate assay and confirmed MBOA-degradation and AMPO-formation in liquid cultures (Classifications in Fig. 4, metabolite data in Fig. S7). MBOA was degraded and AMPO accumulated in cultures of most *Microbacteria* classified as AMPO-formers in the plate assay. Exceptionally, no AMPO was detected for three genomically similar strains (LTA6, LWH12, LWO13). The testing of MBOA metabolisation uncovered four partially related strains (LWH10, LBN7, LWH11 and LWO12) that also accumulated HMPAA, an alternative degradation product of MBOA (Fig. S1). Finally, testing metabolisation of DIMBOA-Glc revealed that also non-AMPO formers degraded DIMBOA-Glc and that most AMPO-forming strains were strong DIMBOA-Glc degraders. An exception of the latter observation was a group of four genomically similar strains that formed AMPO following weak DIMBOA-Glc degradation (LM3X, LMB2, LMX7 and LWO14). For all 39 *Microbacteria,* we further quantified growth in minimal media containing MBOA as sole carbon source, corroborating that AMPO-forming strains have a growth benefit from this trait. Our chemical validation provides a robust basis for comparative genomics of 16 AMPO-forming and 23 AMPO-negative *Microbacteria* strains.

**Figure 4:**
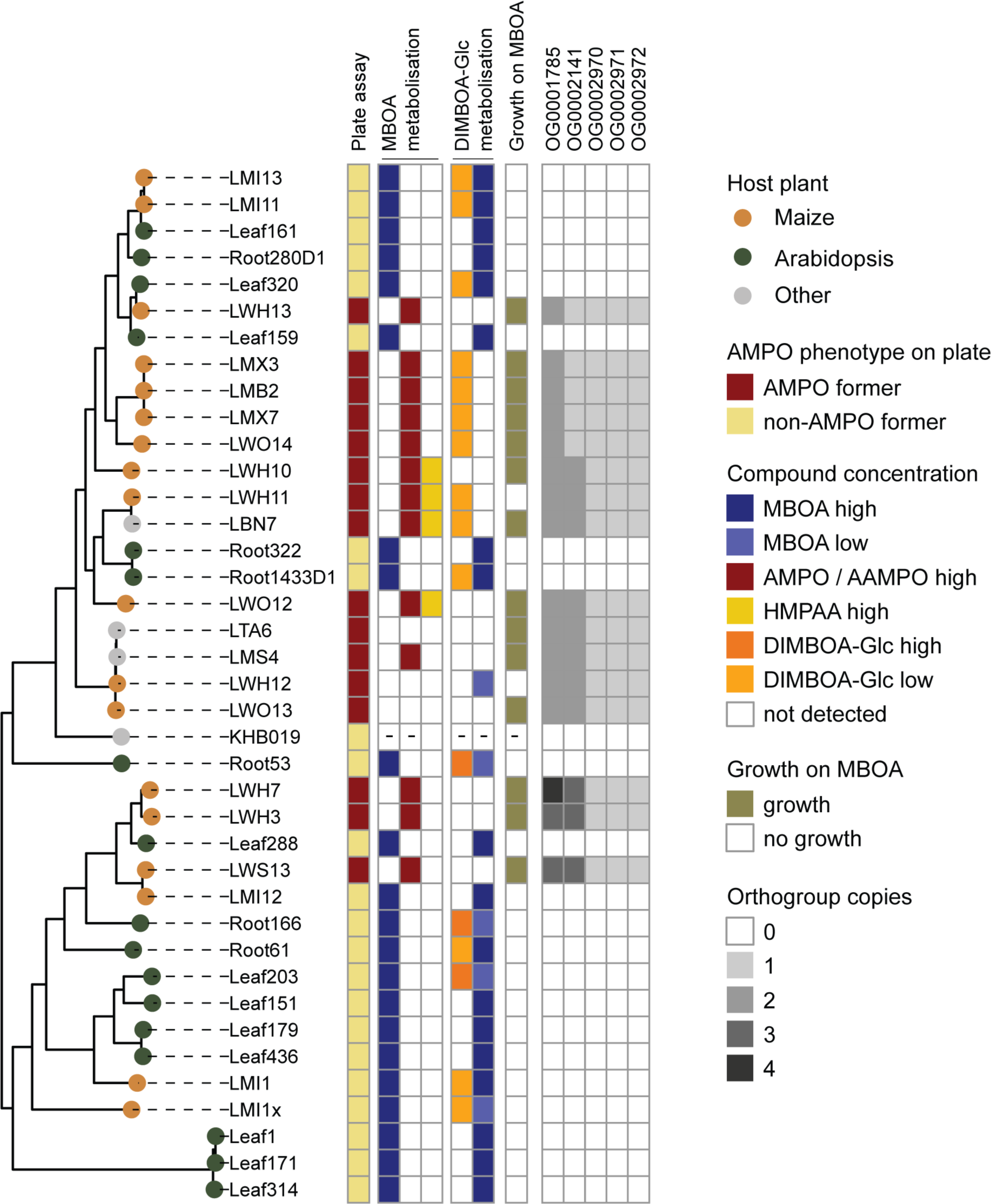
Phenotypic diversity of AMPO-formation in *Microbacteria*. Phylogenetic tree constructed from whole genome alignment of Microbacteria. Tips are coloured by host plant from which the strains were isolated from. The first row shows the AMPO classification (AMPO-former or non-AMPO former) of the strains based on the visual plate assay. The adjacent columns display the qualitative classifications of metabolite analyses (MBOA, AMPO, HMPAA and DIMBOA-Glc) of liquid cultures. The binary scale in column six indicates if the strain grew in minimal medium supplemented with MBOA as a sole carbon source. Mean results of 12 independent replicates grown in two independent runs. Columns seven to eleven report the results from the comparative genomic analysis, representing the copy number of the orthogroups found in each strain.

### Identification of a gene cluster for AMPO-formation in *Microbacteria*

To identify candidate genes for AMPO-formation, we used the AMPO-phenotype of the plate assay and combined three comparative genomic approaches (Supplementary results). The orthogroup method identified 6 candidate genes (Fig. 4, Dataset S2), the kmer approach 17 (Dataset S3) and the transcriptome analysis 108 (Fig. S8, Dataset S4); their overlaps are displayed in Fig. 5a. Mapping the resulting candidates to the genome of LMB2 revealed 15 genes that were located adjacently, pointing to a gene cluster for AMPO-formation (Fig. 5b). This gene cluster contained all 6 genes of the orthogroup analysis, and all 8 genes detected by the kmer approach. Transcripts of the entire gene cluster were significantly upregulated in presence of MBOA, corroborating an active role in AMPO-formation (Fig. S8). We termed this cluster *<u>b</u>*enzo*<u>x</u>*azinoid *<u>d</u>*egradation and named the 15 genes in sequence *bxdA* to *bxdO*. The *bxd* gene cluster encodes 13 enzymes and two transcriptional regulators (Table S1).

**Figure 5:**
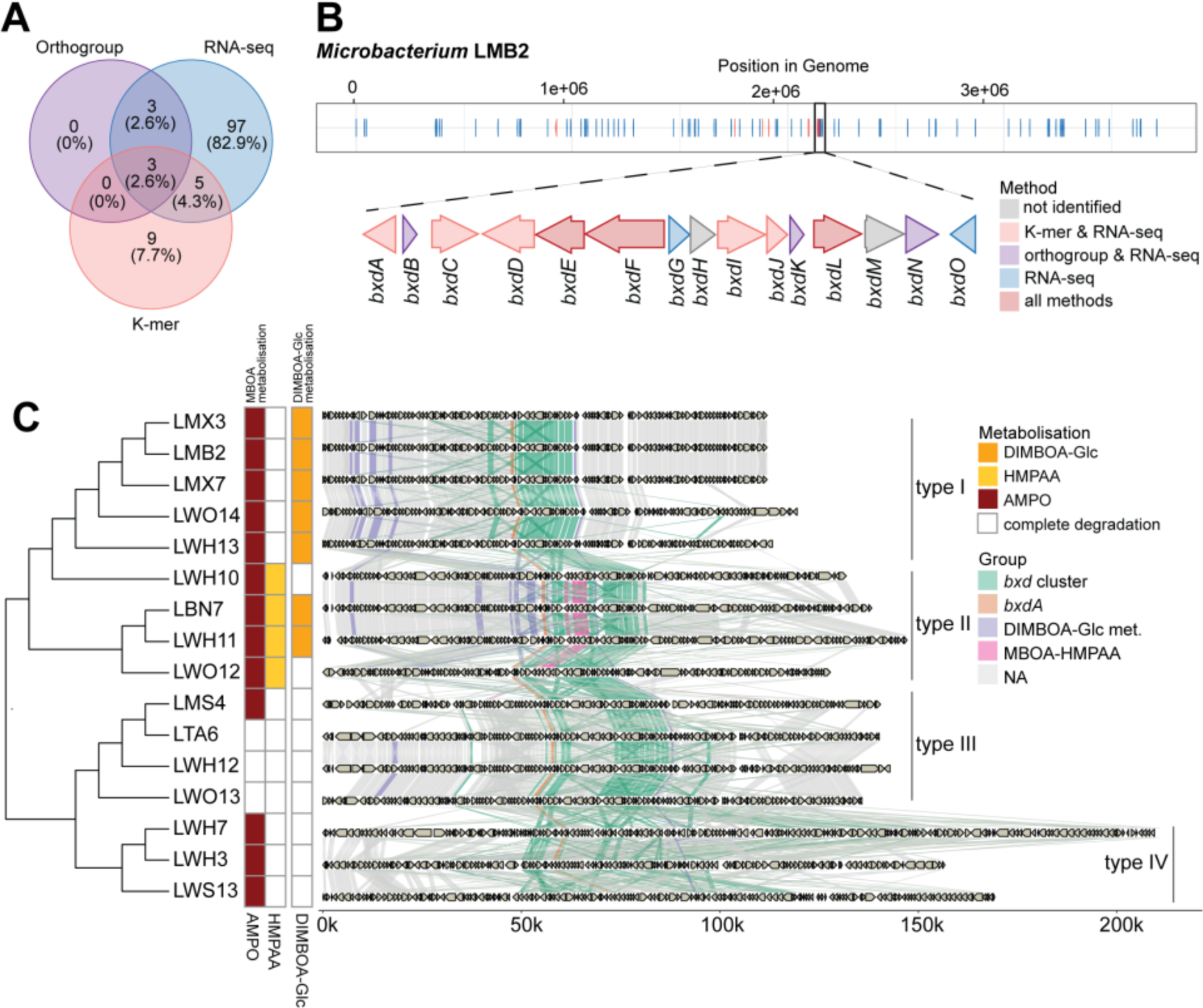
***Bxd* gene cluster in *Microbacteria*. A)** Overlap of three approaches used to identify candidate genes in AMPO formation: orthogroups, kmers and in RNA-seq. **B)** Position of all candidate genes identified with the three approaches in the genome of *Microbacterium* LMB2 with a zoom-in of the *bxd* gene cluster, annotated with its gene architecture including all genes named *bxdA* to *bxdO*. **C)** Across *Microbacteria*, the *bxd* gene cluster consists of four types (type I, type II, type III and type IV) that differ in gene order and content that correspond to their chemical phenotypes.

We performed in-depth analysis of the *bxd* gene cluster on closed long-read genomes of all AMPO-forming *Microbacteria*. High resolution alignments revealed four types of cluster architectures (Fig. 5c). Interestingly, they largely agreed with the different metabolisation phenotypes of the strains (Fig. 4). Gene cluster type I was present in five strains (LMX3, LMB2, LMX7, LWO14 LWH13), all ‘weak DIMBOA-Glc degraders’ that fully degraded MBOA and accumulated AMPO as the only metabolisation product. Type II was found in four strains (LWH10, LWH11, LBN7, LWO12), contained five additional genes in the *bxd* gene cluster, and these strains uniquely formed HMPAA besides accumulating AMPO. Cluster type III, present in four strains (LMS4, LTA6, LWH12, LWO13), corresponded to bacteria that efficiently metabolised DIMBOA-Glc and MBOA without accumulating AMPO. Finally, the cluster type IV, containing many gene duplications, was found in three strains (LWH7, LWH3, LWS13) that all formed AMPO after efficient metabolisation of DIMBOA-Glc or MBOA. This fine-grained genome analysis revealed multiple variants of the *bxd* gene cluster, possibly representing multiple metabolic pathways of benzoxazinoid degradation in *Microbacteria*.

### BxdA converts MBOA to AMPO *in vitro*

To identify the gene(s) responsible for MBOA breakdown and AMPO-formation, we selected the four candidates *bxdA*, *bxdD*, *bxd*G, and *bxdN* based on the functional annotation of their proteins as N-acyl homoserine lactonase family protein (BxdA), aldehyde dehydrogenase family protein (BxdD), VOC family protein (BxdG), and NAD(P)-dependent oxidoreductase (BxdN). We chose heterologous expression in *E. coli* as *Microbacteria* remain genetically unamenable. While neither purified BxdD, BxdG and BxdN nor the empty vector control showed MBOA degrading activity (Fig. S9), purified BxdA degraded MBOA and led to the accumulation of AMPO (Fig. 6a). Hence, the *Microbacteria* gene *bxdA*, encoding a ∼34 kDa protein annotated as an N-acyl homoserine lactonase family protein, has *in vitro* activity to degrade MBOA and form AMPO. We propose that BxdA functions as a lactonase, opening the lactone moiety of MBOA to form 2-amino-5-methoxyphenol (AMP) via the corresponding carbamate (HMPCA) as potential intermediate (Fig. 6b).

**Figure 6:**
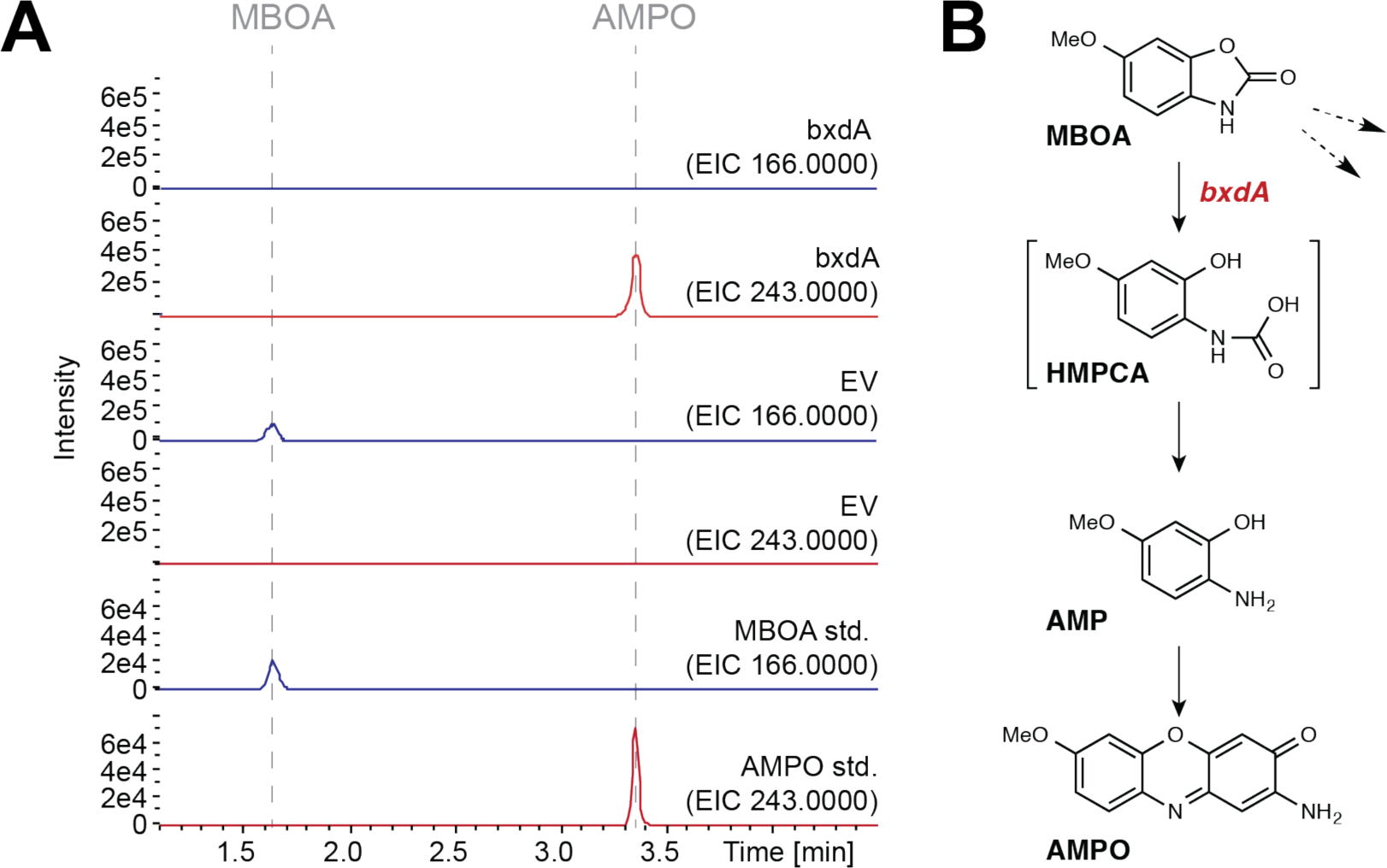
BxdA converts MBOA to AMPO. **A)** Purified recombinant BxdA was incubated with the substrate MBOA and product formation was monitored using high-pressure liquid chromatography-mass spectrometry (HPLC-MS) operated in positive mode (full-scan, EIC = extracted ion chromatogram). An empty vector (EV) control showed no activity. Authentic MBOA and AMPO were used as standards. **B)** Proposed reaction sequence from MBOA to AMPO catalysed by BxdA. The dashed arrows refer to possible alternative MBOA degradation pathways. The potential intermediate HMPCA is proposed but was not confirmed experimentally. HMPCA = (2-hydroxy-4-methoxyphenyl)carbamic acid.

To elucidate whether BxdA is a *Microbacterium*-specific adaptation or a widespread strategy for MBOA degradation, we conducted homology searches (Supplementary results). In brief, besides being present in all our AMPO-forming *Microbacteria, bxdA* was also found in the other strong AMPO-formers (3 gene copies in *Sphingobium*, 1 in *Pseudoarthrobacter*) but not in all other taxa (Fig. S9). In wider homology searches, BxdA was rarely found in other bacteria and unrelated to previously described benzoxazinoid-degrading proteins. Together this highlights the importance of BxdA for AMPO-formation by maize root bacteria and suggests it to be a novel enzyme for microbial metabolisation of benzoxazinoids.

## Discussion

Plants recruit distinct root microbial communities from the soil by exuding bioactive specialised metabolites^5^. Thus, they shape species-specific microbiomes^5^, but the mechanisms are not well understood. Here, we show that many maize root bacteria can metabolise host-exuded benzoxazinoids, the main specialised metabolites of maize. This trait is specific to native root bacteria from maize and is present among taxonomically diverse and abundant members of the root microbiome. Metabolisation of benzoxazinoids was rare in ‘non-host’ Arabidopsis bacteria, i.e., strains isolated from a plant that does not produce benzoxazinoids. Maize bacteria benefitted from metabolising MBOA because they can use it as carbon source in nutrient limiting conditions. Of the different known chemical routes to degrade MBOA (Fig. S1), we have identified *bxdA*, which encodes a novel lactonase enzyme converting MBOA to AMPO. Through *bxdA*, maize root bacteria are metabolically adapted to benzoxazinoid exudates of maize. Below, we discuss metabolic adaptation, the biochemistry of BxdA, and the biological context of these findings.

Root microbes metabolise specialised plant metabolites. For example, Arabidopsis root bacteria degrade host-synthesized triterpenes^28^; several soil microbes metabolise benzoxazinoids^17,37^. Here, we investigated if maize root bacteria are adapted - defined by a heritable trait improving an organism’s fitness - to maize-exuded benzoxazinoids. We found support for this hypothesis by uncovering that benzoxazinoid metabolisation is enriched in maize bacteria, whereas it is missing in non-host bacteria. This differential metabolisation was seen comparing maize^19^ (Fig. 1a) and Arabidopsis strains^36^ (Fig. 2b) and again plating natural root microbiomes of different plant species (Fig. 2a). The finding that AMPO-forming bacteria were less abundant in the wheat root microbiomes may appear surprising but is consistent with the MBOA levels in the rhizosphere of this wheat variety. More than 10x less MBOA (∼5 ng/mL) was found in the rhizosphere of CH Claro compared to maize (∼60 ng/mL)^38^. Hence, we think that the much lower levels of MBOA in the rhizosphere of CH Claro resulted in a much lower selection of AMPO formers.

The fact that roots of benzoxazinoid-deficient *bx1* mutants harboured 50% less AMPO-forming bacteria highlighted a direct link between benzoxazinoid exudation from maize and bacterial MBOA metabolisation (Fig. 1b). Metabolite profiling of *Microbacteria* from maize and Arabidopsis revealed that only maize-derived isolates metabolised benzoxazinoids (Fig. 4), and genomic comparisons uncovered the *bxd* gene cluster, which was only present in AMPO-forming *Microbacteria* (Fig. 5). The key gene *bxdA* was also present in other MBOA-metabolising maize bacteria but not in Arabidopsis-derived bacteria (Fig. S9), which reveals metabolic adaptation of maize root bacteria to host-specialised metabolites at the genomic level. Further research, for instance comparing Arabidopsis and maize root bacteria for metabolisation of specialised compounds of Arabidopsis such as coumarins^26^, is required to broaden this conclusion. Given the high degree of host-species specific microbiomes^5^ and the widespread nature of plant species-specific specialised metabolites, we propose that metabolic adaptation may structure root microbiomes across the plant kingdom.

To complement previous studies^20^, we specifically investigated the genetic basis of benzoxazinoid metabolisation in the native context of root bacteria isolated from benzoxazinoid-exuding maize plants. We focused on MBOA, the most abundant^11^ and most selective^19^ benzoxazinoid in the maize rhizosphere. The phenotypic and genomic screening of maize- and Arabidopsis-derived *Microbacteria* (Fig. 4) permitted the identification of BxdA, an N-acyl homoserine like lactonase. Gene homologs were only found in AMPO-forming *Pseudoarthrobacter* and *Sphingobium* strains from maize but not in Arabidopsis bacteria (Fig. S9). We detected only weak similarity (<43% amino acid level) with known enzymes such as CbaA^31^ or MBL^39^, both involved in metabolisation of benzoxazinoids. Thus, the lactonase BxdA represents a novel enzyme for benzoxazinoid metabolisation pointing to a highly specific adaption restricted to root microbiome members of benzoxazinoid-producing plants.

We confirmed BxdA to catalyse the metabolisation of MBOA to AMPO *in vitro* (Fig. 6a). We chose this approach as *Microbacteria* are genetically unamenable, but future experiments, e.g. with *bxdA* mutants in genera like *Sphingobium*, could allow *in vivo* confirmation. The biochemistry of BxdA is consistent with its annotation as a lactonase that hydrolyses the ester bond of a lactone ring^40^. With MBOA as a substrate, this reaction yields AMP that spontaneously dimerizes to AMPO in the presence of oxygen^24^. Lactonases occur in various bacteria^41^ and typically degrade N-acyl homoserine lactones, which are signalling metabolites of bacterial quorum sensing^42^. This supposedly similar biochemical function opens a range of novel questions, including on the evolutionary origin of BxdA or its impact on quorum sensing that warrant further investigation.

Metabolisation of specialised metabolites has multiple biological consequences. Generally, bacteria aim at detoxification, suppression of other microbes, or utilization as carbon source^43,44^. Our analyses suggested that the bacteria primarily degrade MBOA (Fig. 3c), which is consistent with all AMPO-forming bacteria using MBOA as a carbon source (Fig. 3d & 4). It is conceivable that AMPO is rather formed as a side product of incomplete or inefficient bacterial catabolism of the intermediate AMP. Nevertheless, AMPO-formation will affect the microbial and plant ecology. It confers advantages to AMPO-tolerant bacteria to expand their niche by preferentially suppressing Gram-positive bacteria, which are generally less tolerant to aminophenoxazinones compared to Gram-negative bacteria^19^. Alternatively, AMPO may promote rhizosphere health through its suppressive activity against phytopathogenic fungi^30,45^. Plants, on the other hand, may benefit from recruiting A(M)PO-forming bacteria as they convert (M)BOA to strongly allelopathic compounds that suppress weeds^46^, thereby improving host fitness. Overall, AMPO-forming bacteria contribute to microbiome traits that benefit their host plant.

We had originally discovered benzoxazinoid-dependent and microbe-driven feedbacks on plant performance in controlled conditions^11^ and recently, we show that they also operate in field conditions increasing wheat yield in certain soils^38^. The latter suggest that microbial feedbacks are agriculturally relevant and highlights that plant specialised metabolites present a strong tool for leveraging microbiome functions. In combination, our previous work on tolerance^19^ and the present study demonstrate that metabolic adaptation to plant specialised metabolites are key determinants for root colonisation by bacteria. Regarding possible agricultural applications, our data implies that effective biocontrol or biofertilizer strains should be tolerant and/or metabolically adapted to the specialised metabolites produced by the target crop. Hence, understanding how specific specialized plant metabolites shape and stabilize their microbiomes will be important to harness microbiome functions to improve plant health in sustainable agricultural systems^47^.

## Author contributions

L.T., M.E., and K.S. designed research; L.T. performed microbial plating assays, metabolomic assays, *in vitro* growth assays, performed the transcriptome experiment, and selected candidate genes. M.K. performed comparative genomics and analysed transcriptomic data. M.F. expressed the candidate genes in *E. coli* and tested purified proteins. P.M. performed NMR analyses, P.M, T.Z. and E.K. performed metabolomic analyses. L.R. tested Arabidopsis isolates for AMPO-formation, V.G. cultivated plants, and M.N.D. conducted the anaerobic experiments. S.H., C.B., N.S, C.A.M.R., and T.G.K. provided technical infrastructure and R.B. provided new analytic tools. L.T. and M.K analysed the data and L.T., M.E. and K.S. wrote the manuscript. All authors revised the paper.

## Acknowledgements

We thank Prof. Julia Vorholt (ETH Zurich) and Prof. Paul Schulze-Lefert (MPMI Cologne) for sharing *Microbacteria* strains from the AtSphere collection. Thanks go to Corinne Suter for support with culturing bacteria and plating assays and to Mirco Hecht for supporting metabolomic analysis. Further, we thank Dr. Thomas Roder for the support with the open genome browser, Dr. Pamela Nicholson from the Next-Generation Sequencing Platform in Bern for technical support with sequencing and Dr. Christine Pestalozzi for technical advice. This work was mainly supported by the Interfaculty Research Collaboration “One Health” of the University of Bern. It has also received support by grants of the Austrian Academy of Sciences, the European Union’s Horizon 2020 programme (No. 716823 to C.B.), the European Research Council (No. 189071 to C.R.) and the Swiss National Science Foundation (No. 189071 to C.R.).

## Materials and Methods

### Plating experiment

To assess the number of AMPO-forming colonies on roots, we grew wild-type maize plants and BX-deficient *bx1*(B73) maize, wheat (CH Claro), *Medicago sativa* (Sativa, Rheinau, Switzerland), *Brassica napus* (Botanik Saemereien AG, Pfaeffikon, Switzerland) and *Arabidopsis thaliana* (Col-0) in field soil. The soil was collected in Winter 2019 from the field in Changins^11^. We grew the plants for 7 weeks in a walk-in growth chamber with the following settings: 16:8 light/dark, 26/23 °C, 50 % relative humidity, ∼550 μmol m^-2^s^-1^ light. We fertilized the plants in the following regime: Weeks 1 – 4: 100 mL; 0.2 % Plantactive Typ K (Hauert HBG Duenger AG, Grossaffoltern, Switzerland), 0.0001 % Sequestrene Rapid (Maag, Westland Schweiz GmbH, Dielsdorf, Switzerland); weeks 5 onwards: 200 mL; 0.2 % Plantactive Typ K, 0.02 % Sequestrene Rapid. To account for different need of Arabidopsis growth, all seeds were stratified for three days in the dark at 4 °C and then grown in growth cabinets (Percival, CLF Plant climatics) at 60 % relative humidity, 10 h light at 21 °C and 14 h dark at 18 °C. Arabidopsis were fertilized two times during the experiment by watering with 2/3 water and 1/3 of half-strength Hoagland solution^48^. To harvest the roots, we shake off loose soil and prepared 10 cm long root fragments (corresponding to the depth of -1 to -11 cm in soil) which we then chopped into small pieces with a sterile scalpel. We transferred them into a 50 mL Falcon tube containing 10 mL sterile magnesium chloride buffer supplemented with Tween20 (MgCl^2^Tween, 10 mM MgCl^2^ + 0.05 % Tween, both Sigma-Aldrich, St. Louis, USA). We homogenized the roots with a laboratory blender (Polytron, Kinematica, Luzern, Switzerland; 1 minute at 20’000 rpm) followed by additional vortexing for 15 seconds. For the rhizosphere fraction, we resuspended the pellet from the washing step in 5 mL MgCl^2^Tween. For the soil fraction, we mixed 5 g of soil from the pot with 5 mL MgCl^2^Tween and vortexed it for 15 s.

To quantify bacterial community size, we plated root, rhizosphere, and soil extracts. We serially diluted the extracts and plated 20 µL on 10 % TSB agar (3 g/L tryptic soy broth and 15 g/L agar, both Sigma-Aldrich, St. Louis, USA) plates (12 x 12 cm, Greiner bio-one, Kremsmünster, Austria) containing filter-sterilized cycloheximide (10 mg/L, Sigma-Aldrich, St. Louis, USA) and filter-sterilized DMSO (2 mL/L, Sigma-Aldrich, St. Louis, USA). To spread the drops for counting we tilted the plates and incubated them for 6 days at room temperature. We counted colony-forming units (CFU), multiplied them by the dilution factor and normalized them with the sample’s fresh weight. Before statistical analysis, we transformed CFU counts by log10.

To count the number of AMPO-forming colonies in the extracts, we spread one dilution on a square agar plate containing MBOA. Depending on the plant species and the compartment, we selected a dilution between 1:10-1 and 1:10-4 to reach a colony density which is countable. We spread the 50 μl of the sample with a delta cell spreader on square agar plates with 10% TSB supplemented with filter-sterilized cycloheximide and filter-sterilized MBOA (200 mg/L, Sigma-Aldrich, St. Louis, USA). For 10 days we incubated the plates at room temperature (21 - 25 °C). We photographed the plates and counted the red colonies on the pictures. To get the proportion of AMPO-forming colonies per sample, we divided the count of AMPO-forming colonies by the total CFU.

### Bacterial strains and cultures

Maize root bacteria (i.e., MRB collection)^19^ and Arabidopsis bacteria (i.e., AtSPHERE collection)^36^ were routinely grown on TSA (30 g/L tryptic soy broth and 15 g/L of agar, both Sigma-Aldrich) at 25 °C – 28 °C or TSB liquid medium (30 g/L tryptic soy broth). To screen for AMPO-formation of single isolates, we plated a loop of pure bacterial cultures on TSA plates supplemented with MBOA (200 mg/L) or DMSO (2 mL/L) as control. We incubated the plates for 10 days at room temperature, assessed the phenotype by eye and photographed the plates.

### *In vitro* growth & metabolisation assays

To screen plant root bacteria for their capacity to metabolise benzoxazinoids we deployed the custom, high-throughput, in vitro liquid culture based-growth system reported previously^19,49^. This system makes it possible to culture many bacterial strains in parallel in a replicated manner using many 96-well plates which are handled with a stacker (BioStack 4, Agilent Technologies, Santa Clara, United States), so that the connected plate reader (Synergy H1, Agilent Technologies) records bacterial growth via optical density (OD^600^, absorbance at 600 nm) over time. The assay is set up by inoculating pre-cultures to culture media supplemented with the respective chemical compounds at different concentrations.

Pre-cultures were prepared by transferring isolate colonies with inoculation needles (Greiner bio-one, Kremsmünster, Austria) to 1 mL of liquid 50% TSB (15 g/L tryptic soy broth, Sigma-Aldrich) in 2 mL 96-well deep-well plates (Semadeni, Ostermundigen, Switzerland). These pre-culture growth plates were covered with a Breathe-Easy membrane (Diversified Biotech, Dedham, USA) and grown until stationary phase for 4 days at 28°C and 180 rpm.

Then 4 µL of the pre-cultures were inoculated to 200 µL fresh liquid 50% TSB in 96-well microtiter plates (Corning, Corning, USA) containing the compounds and concentrations to be tested: DIMBOA-Glc, MBOA (500 or 2’500 µM) and BOA (500 μM). These treatments were prepared by mixing their stock solutions into liquid 50% TSB. DIMBOA-Glc was isolated from maize seedlings as described previously^19^ while synthetic MBOA and BOA were commercially available (Sigma-Aldrich). Stock solutions were prepared in the solvent DMSO (Sigma-Aldrich) depending on the solubility of the compounds: DIMBOA-Glc at 500 mM (186.55 mg/mL), MBOA at 606 mM (100 mg/mL) and BOA at 500 mM (67.55 mg/mL). The DMSO concentration was kept constant in each treatment including the control.

All reactions and replicated plates were pipetted using a liquid handling system (Mettler Toledo, Liquidator 96™, Columbus, USA). All plates had lids and were piled up and inserted to a stacker (BioStack 4, Agilent Technologies, Santa Clara, United States), which was connected to a plate reader (Synergy H1, Agilent Technologies, Santa Clara, United States). Using this system, OD^600^ of every culture was recorded every 100 min over 68 h. Prior to each measurement, the plates were shaken for 120 s. In each plate, wells with 50% TSB were included as no bacteria controls (NBC) and in each run one plate containing only media was included to monitor potential contaminations. This procedure applies to the time-series experiment, the Microbacteria screen, the carbon source and BOA assays. To measure MBOA metabolisation over time, we removed plates from the stack after 16 h, 24 h, 44 h, 68 h and 96 h.

For the growth assays with benzoxazinoids as sole carbon source, we followed the same procedure as described above but using minimal media instead of 50% TSB. The minimal media was prepared as described previously^50^ and complemented with defined amounts of stock solutions of either MBOA or DIMBOA-Glc to reach a final concentration of 500 or 2’500 µM. As positive growth controls we grew the bacteria in glucose at different concentrations (500, 2’500 and 30’000 µM) and in 50% TSB.

For the initial metabolite screen of all MRB and the transcriptome experiment, we incubated the plates on a laboratory shaker at 28 °C instead of using the stacker. To avoid evaporation, we sealed the plates with a stripe of Breathe-Easy membrane. We recorded the optical density of the cultures at the end of the experiment in a plate reader (Tecan Infinite M200 multimode microplate reader equipped with monochromator optics, Tecan Group Ltd., Männedorf, Switzerland). The initial metabolite screen ended after 68 h and the transcriptome experiment after 16 h. We exported bacterial growth data from the software (Gen 5, Agilent Technologies, Santa Clara, United States) to excel. We used R statistical software (version 4.0, R core Team, 2016) to analyze growth data. First we calculated the area under the growth curve (x-axis for time and y-axis for OD^600^ AUC) using the function *auc()* from package MESS^51^ and normalized growth in a treatment relative to the control. Such normalized bacterial growth data of a given concentration was statistically assessed (compound vs control) using one-sample t-tests (p-values adjusted for multiple hypothesis testing). Further details on statistical analysis are described below.

### Assessing MBOA metabolisation in anaerobic conditions

To test the requirement of oxygen for AMPO-formation, we performed a metabolisation experiment in anaerobic conditions. As described above, we prepared treatment solutions with 500 μM or 2’500 μM MBOA in 15 mL Falcon tubes. Before the experiment, we pre-incubated the treatments over three days in a sealed jar under an anaerobic environment to remove oxygen from the TSB medium. To start the experiment, we inoculated a loop of bacteria from fresh plates. An anaerobic environment was created for half of the samples with an environment generator according to the manufacturer’s instructions (TRILAB, Jenny Science, Rain, Switzerland). We grew the cultures either under anaerobic or aerobic conditions in an incubator at 28 °C (Memmert, Schwabach, Germany). After 68 h of growth, we measured the optical density of the cultures.

### Metabolite extraction from bacterial cultures

At the end of the experiment, we examined colour changes in the cultures by eye. To fix bacterial cultures, we added 150 μL bacterial cultures to 350 μL of the extraction buffer (100 % Methanol + 0.14 % formic acid) in non-sterile round bottom 96-well plates (Thermo Fisher Scientific, Waltham, USA). We stored the fixed samples with a final concentration of 70 % methanol and 0.1 % formic acid at -80 °C. To reduce the number of samples, we pooled three replicates of the same culture. For the transcriptome experiment (n = 5) and the anaerobic experiment (n = 3), we did not pool samples. We diluted the pooled sample by mixing 50 to 700 μL MeOH 70% + 0.1 % FA. We filtered the cultures through regenerated cellulose membrane filters (CHROMAFIL RC, 0,2 µm, Macherey-Nagel, Düren, Germany) by centrifugation (6’200 rpm for 2 min) to remove bacterial debris. To avoid any residual particles, we centrifuged the extracts at 13’000 rpm for 10 min at 4 °C. We aliquoted the supernatants in glass vials (VWR, Dietikon, Switzerland) and stored the samples for a few days at 20 °C until analysis.

### Profiling benzoxazinoid degradation products in bacterial cultures

Using an Acquity I-Class UHPLC system (Waters, Milford, US) coupled to a Xevo G2-XS QTOF mass spectrometer (Waters, Milford, US) equipped with a LockSpray dual electrospray ion source (Waters, Milford, US) we quantified benzoxazinoids in samples of filtered bacterial cultures. Gradient elution was performed on an Acquity BEH C18 column (2.1 x 100 mm i.d., 1.7 mm particle size (Waters, Milford, US) at 98–50% A over 6 min, 50-100% B over 2 min, holding at 100% B for 2 min, re-equilibrating at 98% A for 2 min, where A = water + 0.1% formic acid and B = acetonitrile + 0.1% formic acid. The flow rate was 0.4 mL/min. The temperature of the column was maintained at 40 °C, and the injection volume was 1 μL. The QTOF MS was operated in sensitivity mode with a positive polarity. The data were acquired over an m/z range of 50–1’200 with scans of 0.1 s at a collision energy of 6 V (low energy) and a collision energy ramp from 10 to 30 V (high energy). The capillary and cone voltages were set to 2 kV and 20 V, respectively. The source temperature was maintained at 140°C, the desolvation temperature was 400 °C at 1’000 L/hr and the cone gas flow was 100 L/hr. Accurate mass measurements (<2 ppm) were obtained by infusing a solution of leucine encephalin at 200 ng/mL at a flow rate of 10 μL/min through the Lockspray probe (Waters, Milford, US). For each expected benzoxazinoid compound, four standards with concentrations of 10, 50, 200, and 400 ng/mL were run together with the samples (DIMBOA-Glc, DIMBOA, HMBOA, MBOA-Glc, MBOA, BOA, AMPO, APO, AAMPO, HMPMA) or 40, 200 ng/mL, 1 and 10 μg/mL for HMPAA and AMP.

### NMR identification of AMPO

To confirm the presence of AMPO in the liquid cultures of *Sphingobium* LSP13 and *Microbacterium* LMB2, we analysed them by ^1^H NMR spectroscopy (Bruker Advance 300, 1H: 300.18 MHz, Bruker Corp., Billerica, MA, USA). Briefly, liquid cultures were centrifuged (20 min, 13’000 rpm) and the supernatants extracted twice with Et^2^O, dried with Na^2^SO^4^ and filtered in a glass funnel with cotton wool. During cultivation a red precipitate formed towards the neck of the Erlenmeyer flasks, i.e. at the edge of the shaking cultures (Fig. S2). This red precipitate left was collected from the Erlenmeyer flasks with acetone. The two extracts were combined, concentrated under reduced pressure, and dried over P^2^O^5^. The ^1^H NMR spectrum of the red residue obtained was recorded in DMSO-*d^6^* and compared to an analytical AMPO standard^23,52^, confirming its presence in our bacterial cultures.

### Phylogenetic tree construction

The phylogenetic tree of all MRB and AtSphere bacteria was computed as described previously^19^. The species tree estimation for *Microbacteria* was obtained from OrthoFinder v. 2.3.8^53^. The 16S trees were reconstructed as follows: First, the 16S sequences were combined into a single FASTA file and then aligned using MAFFT v. 7.475^54^ with default options. The aligned sequences were then used as input to RAxML v. 8.2.12^55^. The multi-threaded version ‘raxmlHPC-PTHREADS’ was used with the options ‘-f a -p 12345 -x 12345 -T 23 -m GTRCAT’ with 1’000 bootstrap replicates. The phylogenetic tree was visualized and annotated in R using the package ggtree^56^.

### Comparative genomics

To find genes that are involved in the transformation of MBOA to AMPO we built an extended collection of 39 *Microbacteria* (MicroE) strains. We selected all *Microbacteria* from maize^19^ (n=18) and from the *AtSphere* collection^36^ isolated from Arabidopsis (n=17) and one strain isolated form clover^57^. Additionally, we selected three strains which we isolated, from root extracts of *Brassica napus* (LBN7), *Triticum aestivum* (LTA6) and *Medicago sativa* (LMS4) due to their red colony phenotype on MBOA plates. For those strains we sequenced the genome by PacBio as described for the MRB collection^19^. The 39 Microbacteria were phenotypically divided into AMPO-forming (n = 16) and AMPO-negative *Microbacteria* (n = 23) strains based on the MBOA plate assay. Two approaches were investigated independently. The first consisted of grouping the genes into orthogroups with OrthoFinder v. 2.3.8^53^ and estimating significant associations between the phenotype and orthogroups by applying Fisher’s Exact Test using the gene trait matching tool in OpenGenomeBrowser^58^. In the second approach, a kmer-similarity search strategy was conducted. The scaffolds of the assemblies were first divided into unique kmers of size 21 base pairs and counted using the tool Kmer Counter v. 3.1.1^59^. The resulting kmer libraries per sample were then merged into a single matrix using custom python scripts. In the next step, the kmers were scored based on their occurrence in AMPO-positive or negative strains. Specifically, the score of a kmer was increased by 1, if the kmer is present in a sample with AMPO-forming phenotype and was decreased by 1 if the kmer is present in a sample with AMPO-negative phenotype. This score can thus be seen as a correlation between genetic sequence and phenotype. The highest scoring kmers were then used to filter genes containing those kmers using custom python scripts. Since this approach relies on exact matches of kmers, the gene sequences containing high-scoring kmers were clustered with a 70% similarity cut-off using vsearch v. 2.17.1^60^. The obtained centroid sequences were then searched with BLAST v. 2.10.0^61^ against a database of all genes from all *Microbacteria* strains using ‘blastn’. The BLAST output was filtered for matches with an e-value < 1e50 which resulted in a list of genes for each centroid sequence. These gene lists were then statistically assessed for their association with the phenotype using Fisher’s Exact Test in R (v. 4.2.1). The p-values were corrected using the Benjamini-Hochberg method.

### Transcriptome analysis

For the transcriptome experiment, bacterial cultures which were grown for 16 h in six individual wells were pooled, and immediately stabilized by the addition of RNAprotect Bacteria Reagent (Qiagen, Hilden, Germany). Bacterial cells were lysed by enzymatic lysis and proteinase K treatment and total RNA was extracted using the RNeasy Mini Kit (Qiagen, Hilden, Germany) with subsequent DNAse treatment using the RapidOut DNA removal kit (Thermo Fisher Scientific, Waltham, USA) following manufacturer’s instructions. The quantity and quality of the purified total RNA were assessed using a Thermo Fisher Scientific Qubit 4.0 fluorometer with the Qubit RNA BR Assay Kit (Thermo Fisher Scientific, Waltham, USA) and an Advanced Analytical Fragment Analyzer System using a Fragment Analyzer RNA Kit (Agilent, Basel, Switzerland), respectively. One hundred ng of input RNA was first depleted of ribosomal RNA using an Illumina Ribo-Zero plus rRNA Depletion Kit (Illumina, San Diego, US) following Illumina’s guidelines. Thereafter cDNA libraries were made using an Illumina TruSeq Stranded total Library Prep Kit (Illumina, San Diego, US) in combination with TruSeq RNA UD Indexes (Illumina, San Diego, US) according to Illumina’s reference guide documentation. Pooled cDNA libraries were sequenced paired end using an Illumina NovaSeq 6000 SP Reagent Kit v1.5 (100 cycles Illumina, San Diego, US) on an Illumina NovaSeq 6000 instrument. The run produced, on average, 14 million reads/sample. The quality of the sequencing run was assessed using Illumina Sequencing Analysis Viewer (Illumina version 2.4.7) and all base call files were demultiplexed and converted into FASTQ files using Illumina bcl2fastq conversion software v2.20. The quality control assessments, generation of libraries and sequencing were conducted by the Next Generation Sequencing Platform, University of Bern.

The quality of the RNA-Seq data was assessed using fastQC v. 0.11.7^62^ and RSeQC v. 4.0.0 2^63^. The reads were mapped to the reference genome using HiSat2 v. 2.2.13^64^. The reference genome of strain LMB2 was prepared before the mapping step as follows: The General Features Format (GFF) file obtained from the assembly was transformed to the Gene Transfer Format (GTF) using AGAT v0.8.0^65^ and subsequently transformed to Browser Extensible Data (BED) format using BEDOPS v. 2.4.39^66^. The HiSat2 index from the reference FASTA file was created using the ‘hisat2-build’ command. FeatureCounts v. 2.0.14^67^ was used to count the number of reads overlapping with each gene as specified in the genome annotation. The Bioconductor package (DESeq2 v1.32.0 5)^68^ was used to test for differential gene expression between the experimental groups. To annotate the genes with Gene Ontology (GO) terms, the genes from the reference assembly were translated to amino acid sequences using the ‘esl-translatè command in HMMER3 v. 3.3.2^69^. Pfam domains were then searched using ‘hmmscan’. GO terms were then mapped to genes and their pfam domains using the pfam2go mapping file (http://current.geneontology.org/ontology/external2go/pfam2go). GO term analysis was performed using the R Bioconductor package TopGO^70^.

### Heterologous expression of candidate genes and protein purification

Plasmids for expression of *bxdA* (N-acyl homoserine lactonase family protein), *bxdD* (aldehyde dehydrogenase family protein), *bxdG* (VOC family protein), and *bxdN* (NAD(P)-dependent oxidoreductase) were ordered from Twist Bioscience. The DNA sequences of the genes were used to generate codon-optimized nucleotide sequences for expression in *E. coli*, applying the default settings. Sequences were introduced to expression plasmid pET28a(+) with *Bam*HI and *Hind*III restriction sites (Twist Bioscience HQ, San Francisco, US). All genes were amplified with Platinum Superfi polymerase II (Thermo Fisher Scientific, Waltham, USA) according to the manufacturer’s instructions by using the following primers for *bxdA* forward AAGTTCTGTTTCAGGGCCCGATGAGTGAGCGTAAAACGGAT and reverse ATGGTCTAGAAAGCTTTACTAAGTTAACAAAATCCCGGC, for *bxdD* forward AAGTTCTGTTTCAGGGCCCGATGGCCATAATGCGGTCCG and reverse ATGGTCTAGAAAGCTTTATTAGGCCACCCAGACAGT, for *bxdG* forward AAGTTCTGTTTCAGGGCCCGATGGCTGACGCTGTACG and reverse ATGGTCTAGAAAGCTTTATTAGCGCTCCGGATGG, for *bxdN* forward AAGTTCTGTTTCAGGGCCCGGTAACTACAGTAGGCTTCTTAG and reverse ATGGTCTAGAAAGCTTTATCAGGACTGGCGGCG and for expression vector pOPINF forward TAATACGACTCACTATAGGG and reverse TAGCCAGAAGTCAGATGCT. Then candidate genes were cloned in the expression vector pOPINF (N-terminal His tag) digested with *Hind*III-HF and *Kpn*I-HF. Cloning was performed with In-Fusion (Takara Bio, Shiga, Japan) according to manufacturer protocol and transformed in chemically competent *E. coli* Top10 (NEB, Ipswich, US) and plated on LB plates (25 g/L Luria-Bertani agar, Carl Roth, Karlsruhe, Germany) supplemented with carbenicillin 100 µg/mL (Sigma-Aldrich, St. Louis, USA). Plasmids were isolated from recombinant colonies and the identity of the inserted sequences was confirmed by Sanger sequencing. Next, the constructs were used to transform chemically competent *E. coli* BL21 (DE3) (NEB, Ipswich, US). Correct uptake of the plasmids was verified through colony PCR with vector specific primers (see above). Positive colonies were inoculated in 5 mL LB with carbenicillin 100 µg/mL and grown overnight at 37 °C, 220 rpm. 100 µL of the preculture were inoculated in 100 mL 2xYT media with carbenicillin 100 µg/mL and incubated at 37°C, 220 rpm until they reached OD^600^ = 0.5-0.6. At this point, cultures were incubated for 15 min at 18 °C, 220 rpm and then induced with IPTG 0.5 mM and incubated at 18 °C, 220 rpm for 16 h. For purification, the cultures were harvested by centrifugation at 3’200 g, 10 min and resuspended in 10 mL of buffer A1 (50 mM Tris-HCl pH 8, 50 mM glycine, 500 mM sodium chloride, 20 mM imidazole, 5% v/v glycerol, pH 8) supplemented with 0.2 mg/mL Lysozyme and EDTA free protease inhibitor cocktail (cOmplete, Roche, Basel, Switzerland) and incubated for 30 min on ice. Cells were disrupted by sonication using a Sonics Vibra Cell at 40% amplitude, 3s ON, 2s OFF, and 2.5 min total time. The crude lysates were centrifuged at 35’000 g for 30 min and the cleared lysates incubated with 200 μL Ni-NTA agarose beads (Takara Bio, Shiga, Japan) for 1h at 4 °C. The beads were then sedimented by centrifugation at 1’000 g for 1 min and washed 4 times with buffer A1 before eluting the proteins with buffer B1 (50 mM Tris-HCl pH 8, 50 mM glycine, 500 mM Sodium Chloride, 500 mM imidazole, 5% v/v glycerol, pH 8). Dialysis and buffer exchange were performed using buffer A4 (20 mM HEPES pH 7.5; 150 mM NaCl) in centrifugal concentrators (Amicon Ultra – 10kDa, Merk Millipore Cork IRL). Proteins were aliquoted in 50 µL and stored at -20 °C. Protein concentration was determined spectrophotometrically at 280 nm on a NanoPhotometer N60 (Implen, Munich, Germany) considering the molecular weight and extinction coefficient. Protein purity and size were checked trough SDS-Page on Novex WedgeWell 12% Tris-Glycine Gel (Invitrogen, Waltham, US). The protein ladder used was Colour Protein Standard Broad Range (NEB, Ipswich, US).

### Enzyme assays and product analysis

All reactions were performed in a total volume of 100 µL, in 25 mM potassium phosphate buffer, pH=7.5 with 5 µg protein. AMPO biosynthetic activity was tested by supplementing the enzyme with 1 mM MBOA (30 mM stock in MeOH, Sigma-Aldrich, St. Louis, USA). In addition, BxdD was supplemented with NADP+ and BxdN with NADP+ and NADPH. Reactions were initiated by protein addition and incubated at 30 °C, 300 rpm for 2 h in the dark. Reactions were quenched by the addition of 100 µL MeOH, incubated on ice for 15 min and then centrifuged at 15’000 g for 15 min. The reactions were filtered through 0.22 μm PTFE syringe filters and then transferred to LC-MS glass vials.

LC-MS analysis was performed on a Dionex UltiMate 3000 UHPLC (Thermo Fisher Scientific, Waltham, USA) equipped with Phenomenex Kinetex XB-C18 column (100 x 2.1 mm, 2.6 µm, 100 Å, column temperature 40 °C) coupled to a Bruker Impact II Ultra-High-Resolution Quadrupole-Time-of-Flight mass spectrometer (Bruker Daltonics) equipped with EVOQ Elite electrospray ionization. Analytical conditions consisted of A: H^2^O + 0.1 % FA and B: ACN, 0.6 mL/min flow with the following gradient: 0-1 min, 15 % B, 1-6 min, 15-35 % B, 6.1-7.5 min, 100 % B, 7.6-10 min, 15 % B. Mass spectrometry data were acquired through ESI with a capillary voltage of 3500 V and end plate offset of 500 V, nebulizer pressure of 2.5 bar with a drying gas flow of 11.0 L/min and a drying temperature of 250 °C. The acquisition was performed at 12 Hz with a mass scan range from 80 to 1’000 m/z. For tandem mass-spectrometry (Ms^2^) collision energy, the stepping option model (from 20 to 50 eV) was used.

### Statistical analysis

We used R version 4.0 (R core Team, 2016) for statistical analysis and visualization of the data. All code used for statistical analysis and graphing is available from https://github.com/PMI-Basel/Thoenen_et_al_BX_metabolisation. For the analysis of bacterial colonisation, we used log transformed data. We checked for normality using Shapiro-Wilk-test. Using t-test or ANOVA we tested for variance. Raw chromatogram data were peak integrated using MassLynx 4.1 (Waters, Milford, US), using defined properties for the reference compounds in the standards. We used the following packages for data analysis and visualizations: Tidyverse^71^, Broom^72^, DECIPHER^73^, DESeq2^68^, emmeans^74^, ggthemes^75^, pheatmap^76^, multcomp^77^, phyloseq^78^, phytools^79^, vegan^80^ in combination with custom functions.

