## supplement for "The lactonase BxdA mediates metabolic adaptation of maize root bacteria to benzoxazinoids"

#### Index

- Supplementary Results
- Supplementary Figures
- Supplementary Tables
- References

### Supplementary Results

#### Identification of gene candidates for AMPO formation

We combined three complementary approaches to narrow down the gene candidates for AMPO formation. First, we compared the genomes using OrthoFinder[1]. Orthogroups are related genes thought to originate from a single gene in the last common ancestor of a clade of species. We found five orthogroups occurring in AMPO-forming strains (Fig. 4). While the orthogroups OG0002970, OG0002971, and OG0002972 contained single copy genes, OG0002141 and OG0001785 were present in varying copy numbers ranging from 1 and 2 to 3 and 4, respectively. Most copies were found in the three Microbacteria (LWS13, LWH3, LWH7). Overall, varying gene copies were found in these five orthogroups of AMPO-forming strains; LMB2 had 6 genes in these 5 orthogroups (Dataset S2).

Second, we screened the genomes for short sequence strings that were associated with AMPO-forming strains using a custom kmer approach (see methods). We identified a total of 377 kmers with a score >=7 across all genomes. Clustering them to the genes and mapping them in all bacteria resulted in 17 gene clusters with significant associations (Fisher's exact test, $p < 0.05$) with the phenotype (Dataset S3).

Third, we performed a transcriptome experiment. We grew the AMPO-forming *Microbacterium* LMB2 for 16 h in MBOA and measured growth, metabolite profiles, and total gene expression relative to its control in DMSO. We assumed that essential transcripts for AMPO formation would be upregulated upon MBOA exposure and should stay active in this short incubation period. We found similar cell numbers of LMB2 (tolerant to MBOA[2]) in both DMSO and MBOA and complete degradation of MBOA and high concentrations of AMPO formed (Fig. S8). The transcript analysis revealed 2.8 % of genes being differentially regulated (108 genes) with 14 down- and 94 upregulated (Dataset S4).

#### Homology searches: *bxdA* is present in AMPO-forming maize root bacteria

After identification of *bxdA,* the N-acyl homoserine lactonase enzyme that initiates the degradation of MBOA, in *Microbacterium* LMB2, we investigated how widespread and similar this gene is within *Microbacteria* and across other bacterial lineages by searching homologs and quantified their similarity on amino acid sequence level using BLASTP. First, among all *Microbacteria* tested in this study and as expected, all AMPO-forming strains possessed homologous *bxdA* proteins with high sequence similarities ranging from 76.25 – 100 % (Fig. S9) while the corresponding gene was missing in AMPO-negative strains or closest protein homologues were of lower than 25% sequence similarity.

Secondly, we searched homologues of *bxdA* among all other, non-*Microbacteria* strains of the maize root bacteria collection, of which genomes were available[2]. Homologues of the lactonase *bxdA* were missing in most genera of the MRB collection. Consistent with the AMPO-forming phenotype, we found similar *bxdA* homologous in *Pseudoarthrobacter* LMD1 and *Sphingobium* LSP13, LMA1 and LMC3 but not in strains that do not form AMPO (Fig. S9). The gene variants of *Pseudoarthrobacter* and *Sphingobium* (3 gene copies) showed amino acid sequences similarities of 78.93% and 58.86-64.87%, respectively). In *Pseudoarthrobacter* we found the *bxd* gene cluster (except the aldehyde dehydrogenase family protein) organized like type I in LMB2 and this was consistent with the chemical phenotype of *Pseudoarthrobacter* LMD1 of degrading MBOA and forming only AMPO. In *Sphingobium* we found three copies of *bxdA*. In the proximities of the lactonase, we identified several genes of the original *bxd* gene cluster of LMB2 including the M24 family metallopeptidase, NAD(P)-dependent oxidoreductase, VOC family protein and two copies of the MFS transporter. In summary, we only find homologous *bxdA* genes in AMPO-forming strains and the similarity of genetic architecture of the *bxd* gene clusters suggests conservation of this genetic element of benzoxazinoid metabolisation.

Third, we searched homologues beyond our collection of maize root bacteria and blasted *bxdA* against the NCBI database[3]. Most similar *bxdA* genes were identified in bacteria of the Micrococcaceae family, e.g., in an *Arthrobacter* sp. (77.89 % amino acid sequence similarity) or a *Leucobacter* sp. (76.17 %; Fig. S9, Dataset S5). We also identified *bxdA*-like genes in more distantly related bacteria, specifically in members of the Burkholderiaceae family like *Paraburkholderia* sp. (63.82%) or in the Pseudomonadaceae, namely in *Pseudomonas poae* (59.67%). The fact that we only find homologous genes with < 80% sequence similarity and that they are present only in a few different families, indicates that this gene rarely found among bacteria represented in the searched database (Dataset S5).

Finally, we compared the *bxdA* from *Microbacterium* LMB2 with proteins previously reported to act in the metabolization of benzoxazinoids. For instance, a metaldependent hydrolase CbaA was identified in the bacterium *Pigmentiphaga,* an enzyme catalysing the degradation of a derivate of a benzoxazinoid to the corresponding aminophenoxazinone[4]. The metallo-ß-lactamases (mbl) of the fungus *Fusarium pseudograminearum* were found to degrade a benzoxazinoid[5]. The *bxdA* gene only shared very low 42.58% and 30.11% sequence similarity to *cbaA* and *mbl*, respectively. This is consistent with the different annotated enzymatic functions of *bxdA*, *cbaA* and *mbl*, possibly acting in different pathways of benzoxazinoid degradation.

### Supplementary Figures

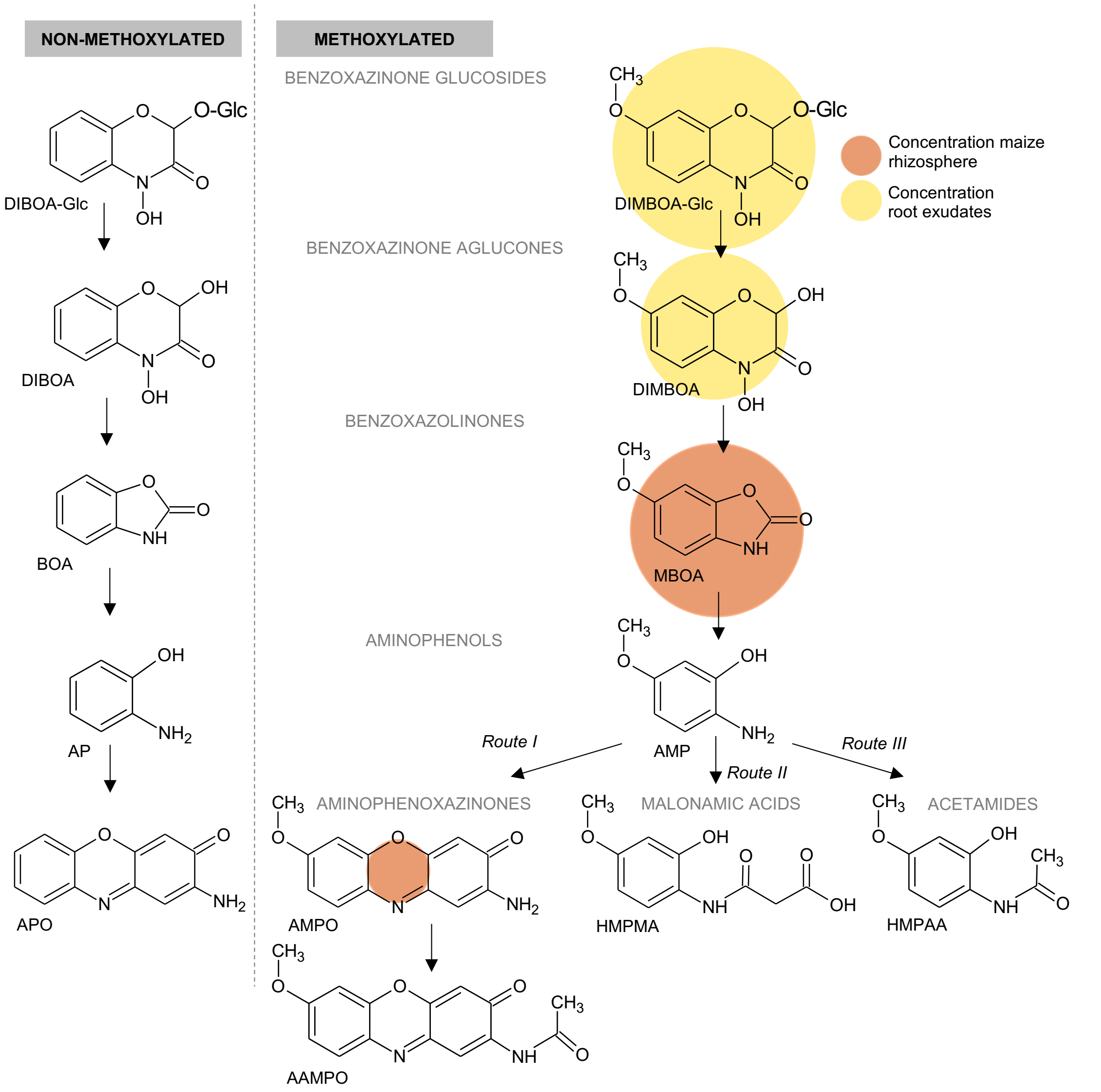


| Abbreviation | Full name | Class | Mass [g/mol] | Formula | Conc. rhizosphere [µg/kg FW]* | Conc. root exudates [µg/kg FW]* |
|---|---|---|---|---|---|---|
| DIMBOA-Glc | 4-hydroxy-7-methoxy-2-[3,4,5-trihydroxy-6-(hydroxymethyl)oxan-2-yl] oxy-1,4-benzoxazin-3-one | Benzoxazinone glucoside | 373.31 | C15H19NO10 | 0.00±0.00 | 0.1±0.02 |
| DIBOA-Glc | 4-hydroxy-2-[3,4,5-trihydroxy-6-(hydroxymethyl)oxan-2-yl]oxy-1,4-benzoxazin-3-one | Benzoxazinone glucoside | 343.29 | C14H17NO9 | NA | NA |
| DIMBOA | 2,4-dihydroxy-7-methoxy-1,4-benzoxazin-3-one | Benzoxazinone aglucone | 211.17 | C9H9NO5 | 0.57±0.34 | 0.05±0.02 |
| DIBOA | 2,4-dihydroxy-1,4-benzoxazin-3-one | Benzoxazinone aglucone | 181.15 | C8H7NO4 | NA | NA |
| MBOA | 6-methoxybenzoxazolin-2(3H)-one | Benzoxazolinone | 165.15 | C8H7NO3 | 5.62±1.39 | 0.002±0.000 |
| BOA | Benzoxazolin-2(3H)-one | Benzoxazolinone | | | ND | ND |
| AMP | 2-amino-5-methoxyphenol | Aminophenol | 139.15 | C7H9NO2 | ND | ND |
| HMPAA | N-(2-hydroxy-4-methoxyphenyl)acetamide | Acetamide | 181.19 | C9H11NO3 | ND | ND |
| HMPMA | N-[2-hydroxy-4methoxyphenyl]malonamic acid | Malonamic acid | 225.2 | C10H11NO5 | ND | ND |
| AMPO | 2-amino-7-methoxy-phenoxazin-3-one | Aminophenoxazine | 242.23 | C13H10N2O3 | 0.39±0.10 | ND |
| AAMPO | 2-acetylamino-7-methoxy-phenoxazin-3-one | Aminophenoxazine | 284.27 | C15H12N2O4 | ND | ND |

*Measurements from Hu et al. 2018, ND = not detected

**Figure S1: Benzoxazinoid metabolites produced by maize and degradation pathways in soil reported in literature.** Bubble size represent the amounts of the compounds measured in root exudates (yellow) and in the rhizosphere (orange). Table lists the full chemical name, the compound class, the molar mass, the chemical formula and the concentrations ± standard deviation based on the measurements from Hu et al. 2018, ND = not detected.

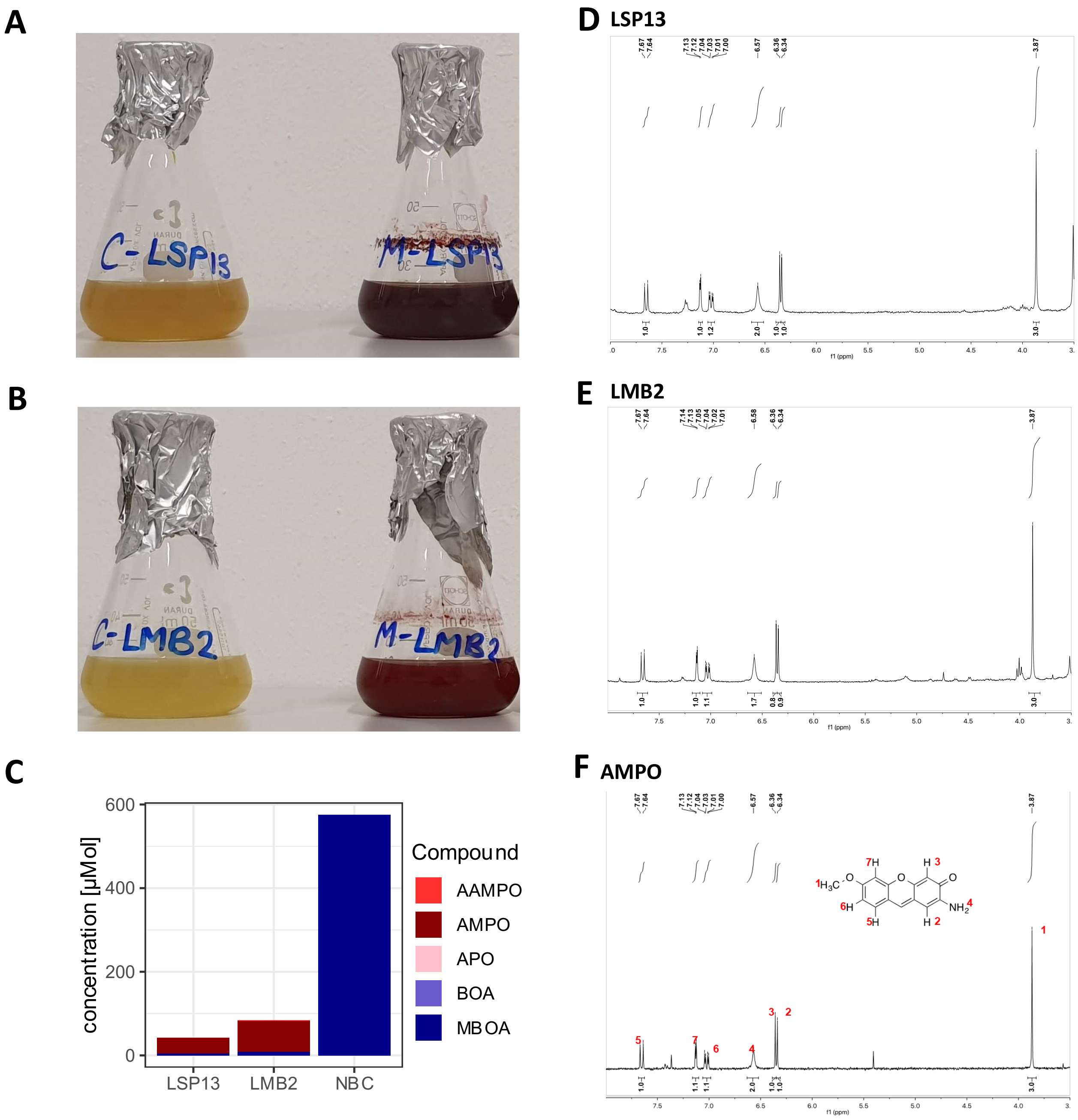


**Figure S2: AMPO phenotype and confirmation of AMPO formation by NMR.** **A)** Pictures of pure cultures in DMSO (left) and MBOA (right) of AMPO-forming strains *Sphingobium* LSP13 and **B)** *Microbacterium* LMB2. **C)** Metabolite profiles of LSP13 and LMB2 grown in MBOA for 68 hours. **D)** NMR spectra of the red precipitate purified from cultures grown in MBOA-supplemented liquid medium for 68 h of LSP13 and **E)** LMB2 and **F)** a pure AMPO sample. The pattern of peaks in the red precipitate extracted from bacterial cultures matches with pure AMPO.

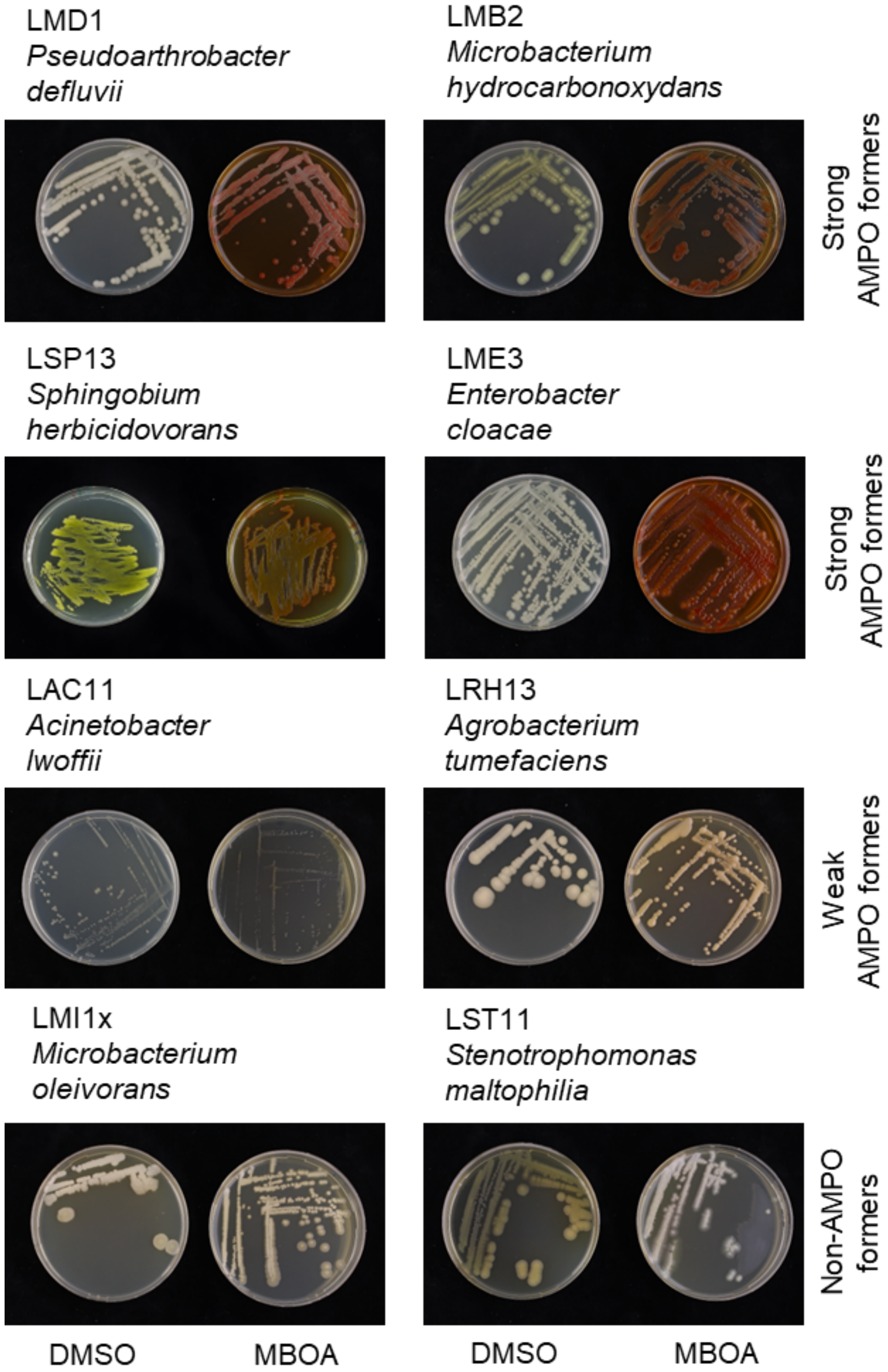


**Figure S3: Rapid screening method for AMPO-formation.** AMPO-forming strains from maize root bacteria strain collection plated on medium containing DMSO (left) or MBOA (right) and incubated for 10 days. Strong AMPO producers form a strong red colour on MBOA medium while weak AMPO producers form less. As a negative control two non-AMPO-forming strains are shown.

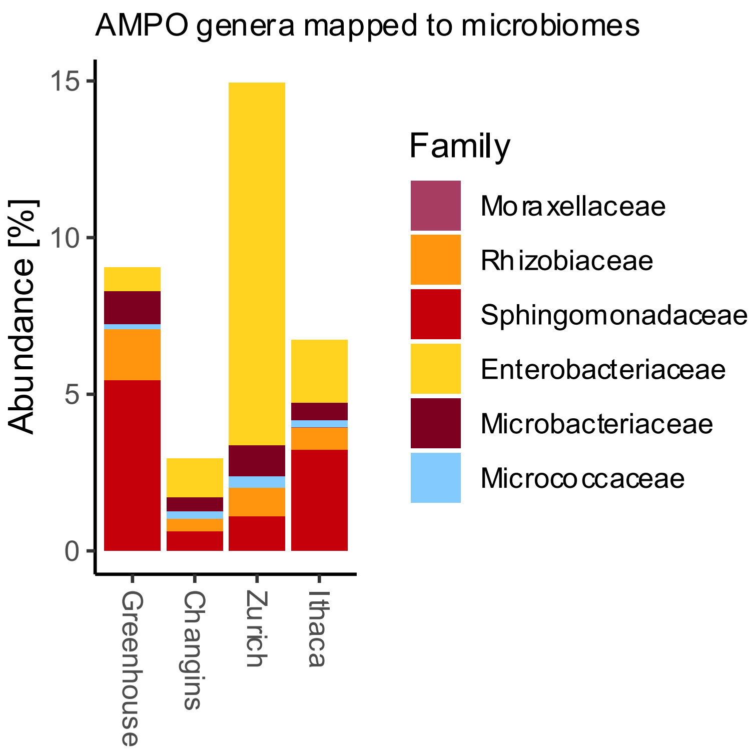


**Figure S4: AMPO-forming colonies are abundant microbiome members on BX-producing maize roots.** Cumulative relative abundance of taxonomic units in field soil represented by AMPO-forming isolates. Datasets from greenhouse experiment with field soil and fields in Switzerland (Changins and Zurich) and the US (Ithaca), Hu et al. 2018 and Cadot et al. 2021 were used for this analysis.

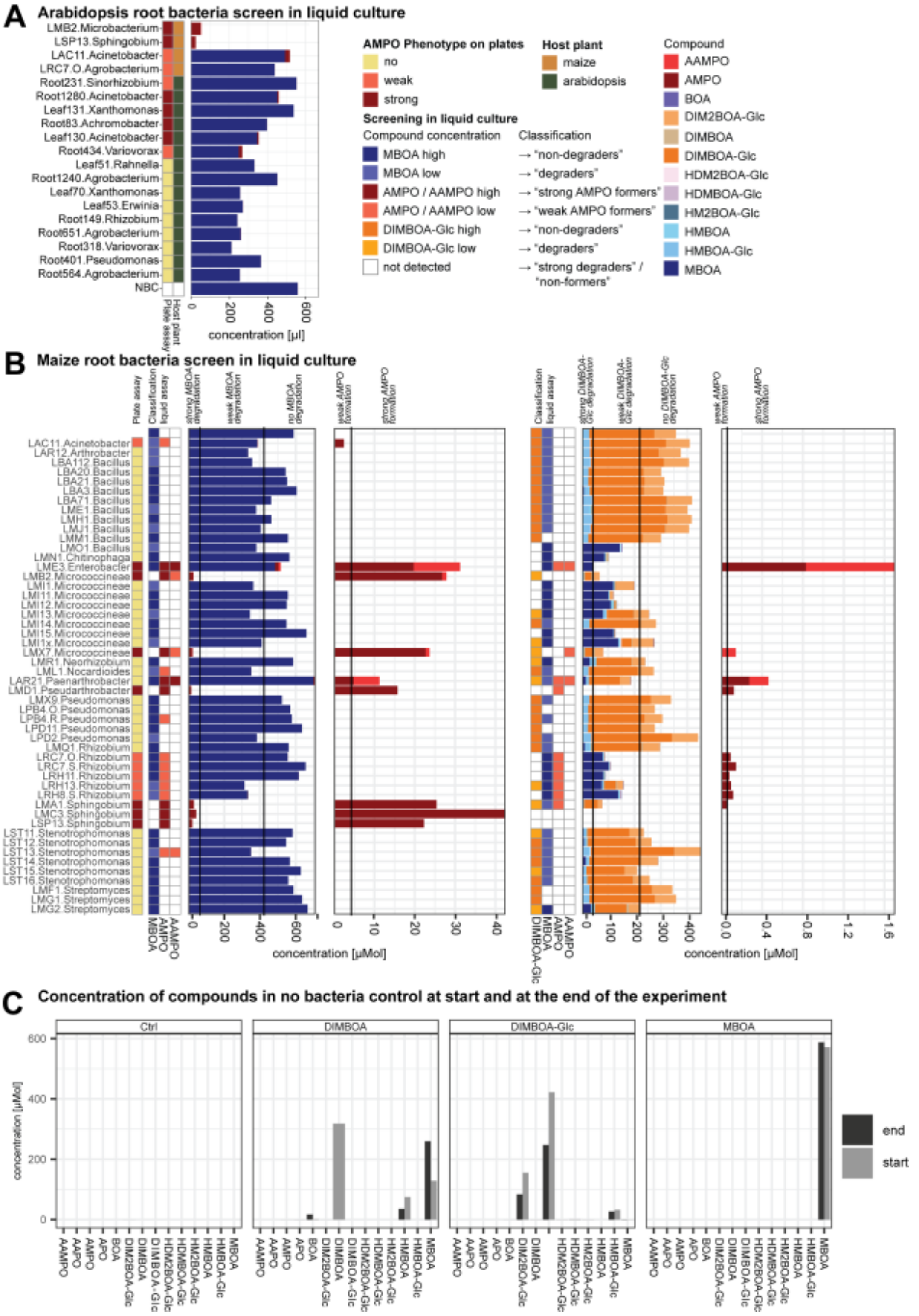


**Figure S5. Complete metabolisation of benzoxazinoids by Arabidopsis bacteria and maize root bacteria. A)** Metabolisation products represented in stacked bargraphs form single strains from MRB strain collection supplemented with DIMBOA-Glc or MBOA. **B)** Only AMPO and AAMPO formation in the tested conditions. **C)** Concentration of DIMBOA-Glc and MBOA in treatment solutions at the start of the experiment (T0) and at the end (NBC). **D)** MBOA and BOA and metabolisation products from selected MRB. **E)** MBOA metabolisation by AtSphere bacteria. Strains with weak colour change on plates, negative and AMPO-forming MRB were compared. All measurements were made from three independently grown samples which were pooled in equal ratios prior to metabolite analysis.

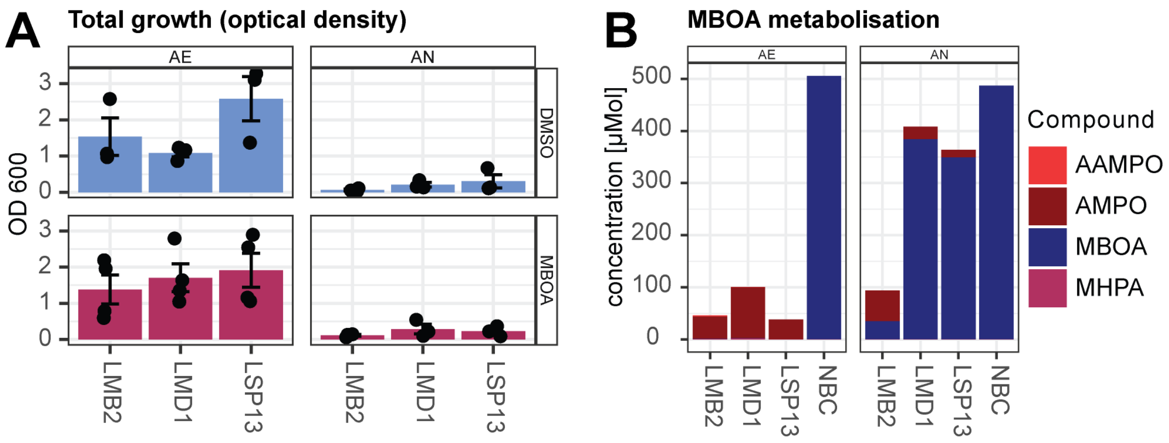


**Figure S6: MBOA metabolisation by three selected strains in aerobic (AE) and anaerobic (AN) conditions. A)** Metabolisation profile of strains grown in MBOA for 68 h both conditions. Replicates are shown in single bars. Concentrations shown in µM. **B)** Bacterial growth of cultures after 68 hours (OD600) in DMSO and MBOA treatment in aerobic and anaerobic condition. **C)** Pictures of cultures at the end of the experiment.

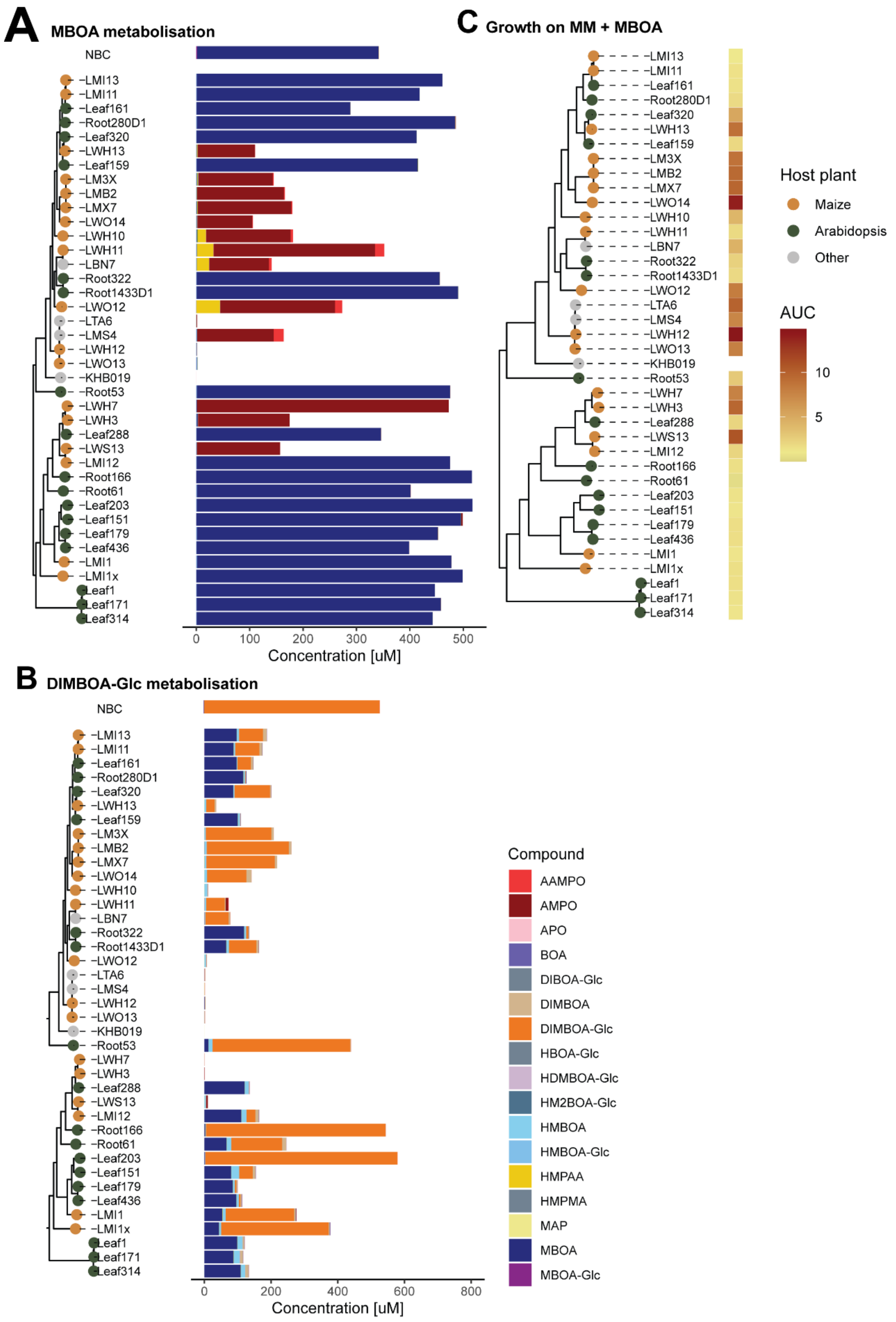


**Figure S7: Benzoxazinoid metabolisation by *Microbacteria***: Phylogenetic tree annotated with metabolite profiles of **A)** MBOA and **B)** DIMBOA-Glc as bar graphs. **C)** Total growth (AUC, area under the curve of growth curve over 68 h) in minimal medium with MBOA as a sole carbon source. Represented values are mean values from 12 independently grown samples in two independent experiments.

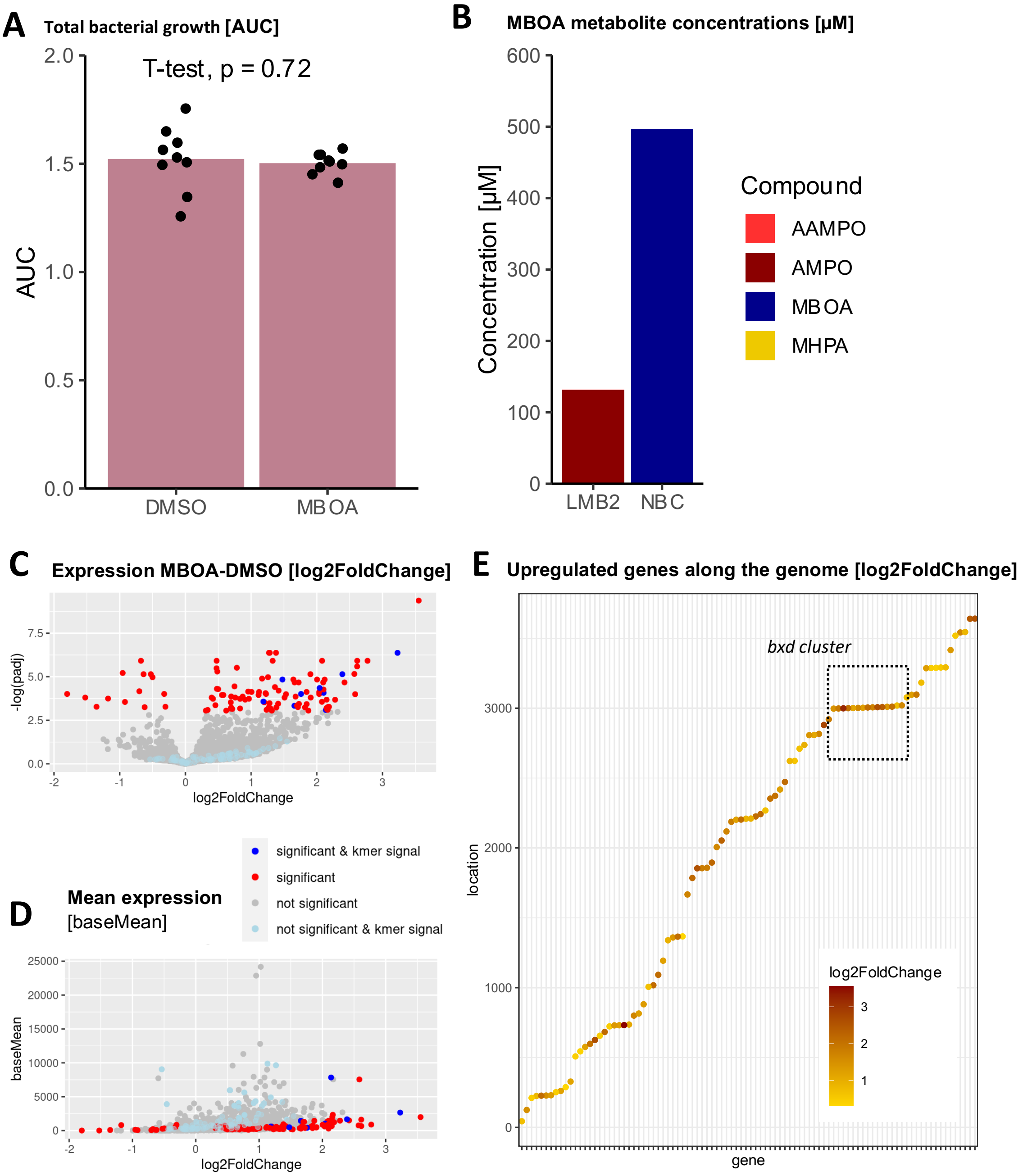


**Figure S8: Transcriptomic experiment of *Microbacterium* LMB2. A)** Total growth of cultures assessed by optical density (OD600) measurements calculated to area under the curve (AUC). **B)** MBOA metabolisation profile of LMB2 and the negative control without bacteria (NBC). All measurements were made from six independently grown samples which were pooled in equal ratios prior to metabolite analysis. **C)** Vulcano plot representing differentially regulated genes, a dotplot representing the expression of the differentially regulated genes and a VennDiagramm showing the overlap of genes differentially expressed in both strains. **D)** A visualization of the differentially expressed genes over the whole genome, highlighting the *bxd* gene cluster in LMB2.

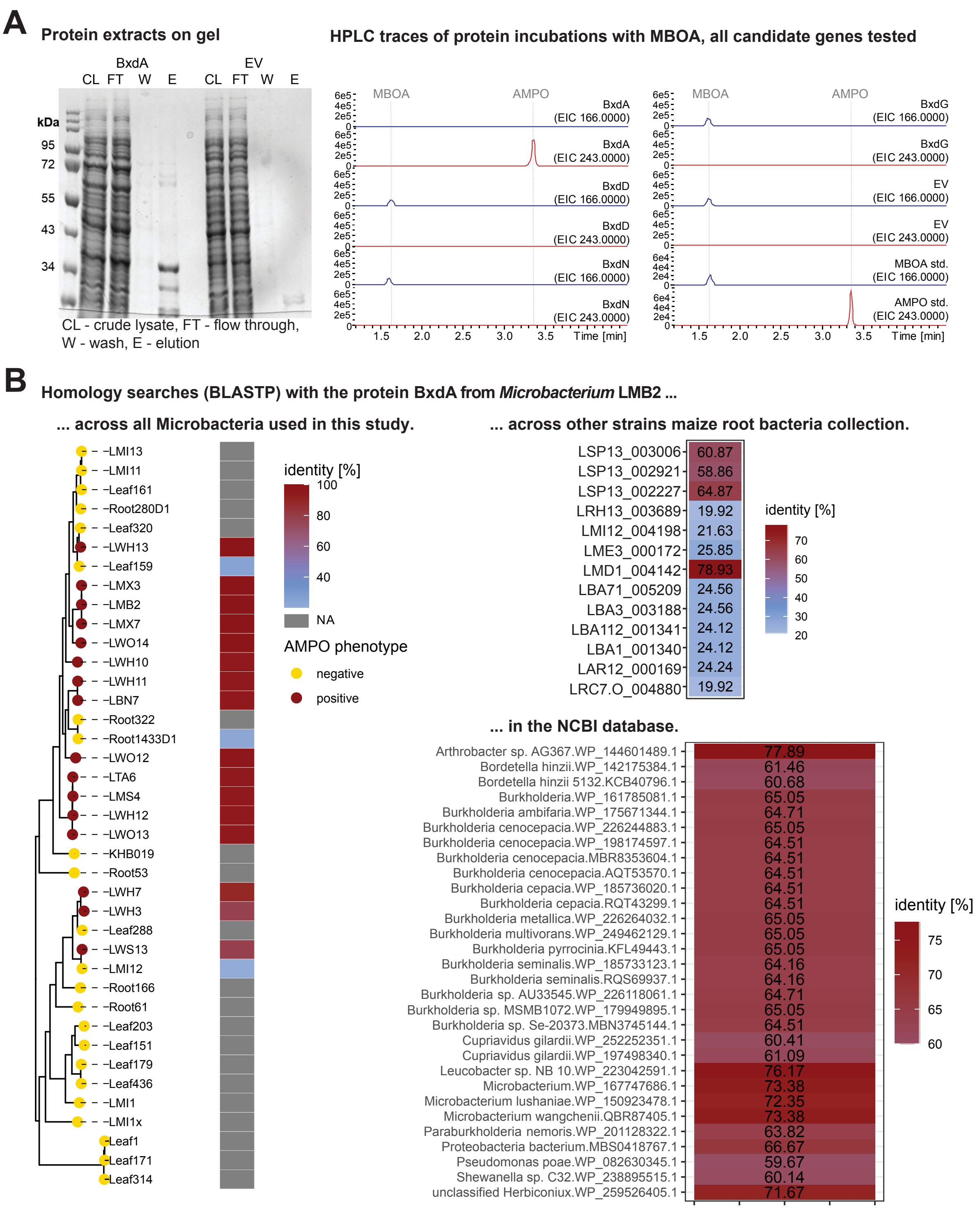


**Figure S9: BxdA converts MBOA to AMPO and is specific to maize bacteria. A)** Gel of purified proteins from *E. coli* cultures with *bxdA* and the empty vector (EV) constructs. Purified recombinant proteins of *E. coli* cultures expressing *bxdA*, *bxdD*, *bxdG*, *bxdN* or the EV construct were incubated with the substrate MBOA and product formation was monitored with high pressure liquid chromatography-mass spectrometry (HPLC-MS) operated in positive mode (full-scan, EIC = extracted ion chromatogram). The EV control, BxdD, BxdG and BxdN showed no activity. Authentic MBOA and AMPO were used as standards. **B)** Homology searches with the protein BxdA of the Microbacterium strain LMB2 (i) across all Microbacteria used in this study, (ii) across all strains of our MRB collection and (iii) against the NCBI database. BLASTP outputs report the % protein similarity. The top 30 hits from the NCBI database are reported (accessed September 2022).

### Supplementary Tables

**Table S1: List of genes present in the *bxd* gene cluster**

| Gene | Annotation | Type |
|---|---|---|
| ***bxdA*** | N-acyl homoserine lactonase family protein | Enzyme |
| ***bxdB*** | RidA family protein | Enzyme |
| ***bxdC*** | acyl-CoA dehydrogenase family protein | Enzyme |
| ***bxdD*** | aldehyde dehydrogenase family protein | Enzyme |
| ***bxdE*** | thiamine pyrophosphate-dependent enzyme | Enzyme |
| ***bxdF*** | 2-oxo acid dehydrogenase subunit E2 | Enzyme |
| ***bxd*** | VOC family protein | Enzyme |
| ***bxdH*** | GntR family transcriptional regulator | Transcriptional regulator |
| ***bxdI*** | acyl-CoA dehydrogenase family protein | Enzyme |
| ***bxdJ*** | flavin reductase | Enzyme |
| ***bxdK*** | RidA family protein | Enzyme |
| ***bxdL*** | M24 family metallopeptidase | Enzyme |
| ***bxdM*** | LacI family DNA-binding transcriptional regulator | Transcriptional regulator |
| ***bxdN*** | NAD(P)-dependent oxidoreductase | Enzyme |
| ***bxdO*** | NADPH-dependent F420 reductase | Enzyme |

#### Supplementary Datasets

**Dataset S1**: Table listing all bacterial strains used for this study including the three Microbacteria isolated and sequenced in this study for extended Microbacteria collection (MicroE), maize root bacteria (MRB) and Arabidopsis bacteria (AtSphere).

**Dataset S2:** Excel file listing all the results of the OrthoFinder approach for all orthogroups across the genome across the Microbacteria.

**Dataset S3:** Table listing the kmers with the highest scores across the Microbacteria.

**DatasetS4:** The file reporting the expression, the differential change between the treatments and the statistics of all the genes in the LMB2 genome.

**DatasetS5:** Excel file including the results of the blast of bxdA to the NCBI database.
